# HDAC1 SUMOylation promotes Argonaute directed transcriptional silencing in *C. elegans*

**DOI:** 10.1101/2020.08.17.254466

**Authors:** Heesun Kim, Yue-He Ding, Gangming Zhang, Yong-Hong Yan, Darryl Conte, Meng-Qiu Dong, Craig C. Mello

## Abstract

Eukaryotic cells use guided search to coordinately control dispersed genetic elements. The transitive effectors of these mechanisms, Argonaute proteins and their small-RNA co-factors, engage nascent RNAs and chromatin-associated proteins to direct transcriptional silencing. The small ubiquitin-like modifier (SUMO) has been shown to promote the induction and maintenance of silent chromatin (called heterochromatin) in yeast, plants, and animals. Here we show that Argonaute-directed transcriptional silencing in *C. elegans* requires SUMOylation of the type 1 histone deacetylase HDA-1. SUMOylation of HDA-1 promotes interactions with components of the nucleosome remodeling and deacetylase (NuRD) complex and with the nuclear Argonaute HRDE-1/WAGO-9. Our findings suggest how HDAC1 SUMOylation promotes the association of HDAC and other chromatin remodeling factors with a nuclear Argonaute in order to initiate de novo heterochromatin silencing.

## Introduction

Argonautes are an ancient family of proteins that utilize short nucleic-acid guides (usually composed of 20-30 nts of RNA) to find and regulate cognate RNAs (Reviewed in Meister, 2013). Argonaute/small-RNA pathways are linked to chromatin-mediated gene regulation in diverse eukaryotes, including plants, protozoans, fungi, and a variety of animals (Reviewed in Martienssen and Moazed, 2015). Connections between Argonautes and chromatin are best understood from studies in the fission yeast *S. pombe*, where an RNA-induced transcriptional silencing (RITS) complex—composed of an Argonaute, Ago1, a novel protein, Tas3, and a heterochromatin protein 1 (HP1) homolog, Chp1—maintains and expands heterochromatin (Verdel et al., 2004). The Chp1 protein binds H3K9me3 through its conserved chromodomain (Partridge et al., 2000; Partridge et al., 2002) and is thought to anchor Ago1 within heterochromatin, where it is poised to engage nascent RNA transcripts (Holoch and Moazed, 2015b). The low-level transcription of heterochromatin is thought to create a platform for propagating small-RNA amplification and heterochromatin maintenance (Reviewed in Holoch and Moazed, 2015a).

In yeast, a protein complex termed SHREC (Snf2/Hdac-containing Repressor complex) has been linked to both the establishment and maintenance of heterochromatin (Sugiyama et al., 2007). SHREC contains two enzymatic activities a histone deacetylase homologous to type 1 HDACs and an ATP-dependent chromatin remodeler homologous to the vertebrate proteins Mi-2 and CHD3. SHREC also contains a Krüppel-type C2H2 zinc finger protein. In all these respects SHREC shares core components found in related eukaryotic Nucleosome Remodeling and Deacetylase Complexes, NuRD complexes, that often differ from each other with regard to the specific Krüppel-type C2H2 zinc finger and other accessory factors they employ (Denslow and Wade, 2007; Torchy et al., 2015). Studies in yeast suggest that SHREC is required broadly for heterochromatin formation and transcriptional silencing (Job et al., 2016; Motamedi et al., 2008; Sugiyama et al., 2007), and similarly NuRD complexes have been shown to function broadly in both developmental gene regulation and transposon silencing (Ecco et al., 2017; Feschotte and Gilbert, 2012; Ho and Crabtree, 2010). These factors are thought to play a key role in converting chromatin from an active to a silent state, and have been shown to be recruited to targets both through sequence-independent interactions with either chromatin or methylated DNA, and through sequence-specific interactions mediated by their Krüppel-type C2H2 zinc finger co-factors and other associated DNA-binding factors (Ecco et al., 2017; Lupo et al., 2013).

The post-translational modification of heterochromatin factors by the small ubiquitin-like protein SUMO has been implicated at several different steps in the establishment and maintenance of transcriptional gene silencing and has been linked to Piwi-Argonaute mediated silencing (Reviewed in Ninova et al., 2019). The addition of SUMO (i.e., SUMOylation) requires a highly conserved E2 SUMO-conjugating enzyme, UBC9, which interacts with substrate-specific co-factor (E3) enzymes to covalently attach SUMO to lysines in target proteins (Johnson, 2004). SUMOylation can have diverse effects, and is not primarily associated with the turnover of its targets, but rather is often associated with changes in protein interactions, especially interactions with proteins that contain SUMO-interacting motifs (SIMs) (Reviewed in Kerscher, 2007). In mammals, SUMOylation has been linked to transposon silencing mediated by an expanded group of Krüppel-Associated Box (KRAB) domain factors (Ecco et al., 2017; Ivanov et al., 2007), which are thought to bind transposon DNA through sequence-specific interactions mediated by their C2H2 zinc finger domains and to further promote recruitment of a NuRD complex and a histone SETDB methyl transferase to silence their targets (Ivanov et al., 2007).

Here we identify a connection between the SUMO pathway and Argonaute mediated-transcriptional silencing in *C. elegans*. We show that SUMOylation of a type-1 HDAC, HDA-1, on C-terminal lysines activates its histone deacetylase activity and is required for Piwi-mediated transcriptional silencing in *C. elegans*. SUMOylation of HDA-1 promotes its association with a Krüppel-type zinc finger protein MEP-1 and with other conserved components of a *C. elegans* NuRD complex. HDA-1 SUMOylation also promotes association of HDA-1 with a nuclear Argonaute HRDE-1/WAGO-9 as well as histone demethylase SPR-5, and the SetDB-related histone methyltransferase MET-2. Our findings suggest how SUMOylation of HDAC1 promotes the recruitment and assembly of an Argonaute-guided chromatin remodeling complex to orchestrates de-novo gene silencing in the *C. elegans* germline.

## Results

### The SUMO and HDAC pathways promote piRNA silencing

To identify additional components of the transcriptional-silencing arm of the piRNA pathway, we conducted an RNAi-based genetic screen of known chromatin factors and modifiers. We utilized a piRNA sensor assay, in which a bright easily scored *gfp::csr-1* fusion transgene is 100% silenced in wild-type germlines, but 100% active in the germlines of piRNA pathway mutants such as *prg-1(tm872)* and *hrde-1/wago-9(ne4769)* (Figure 1A)(Seth et al., 2018). We found that even the partial inactivation of piRNA components by RNAi was sufficient to activate GFP::CSR-1 expression in a percentage of exposed animals (Figure 1B and Table S1).

**Figure 1.**
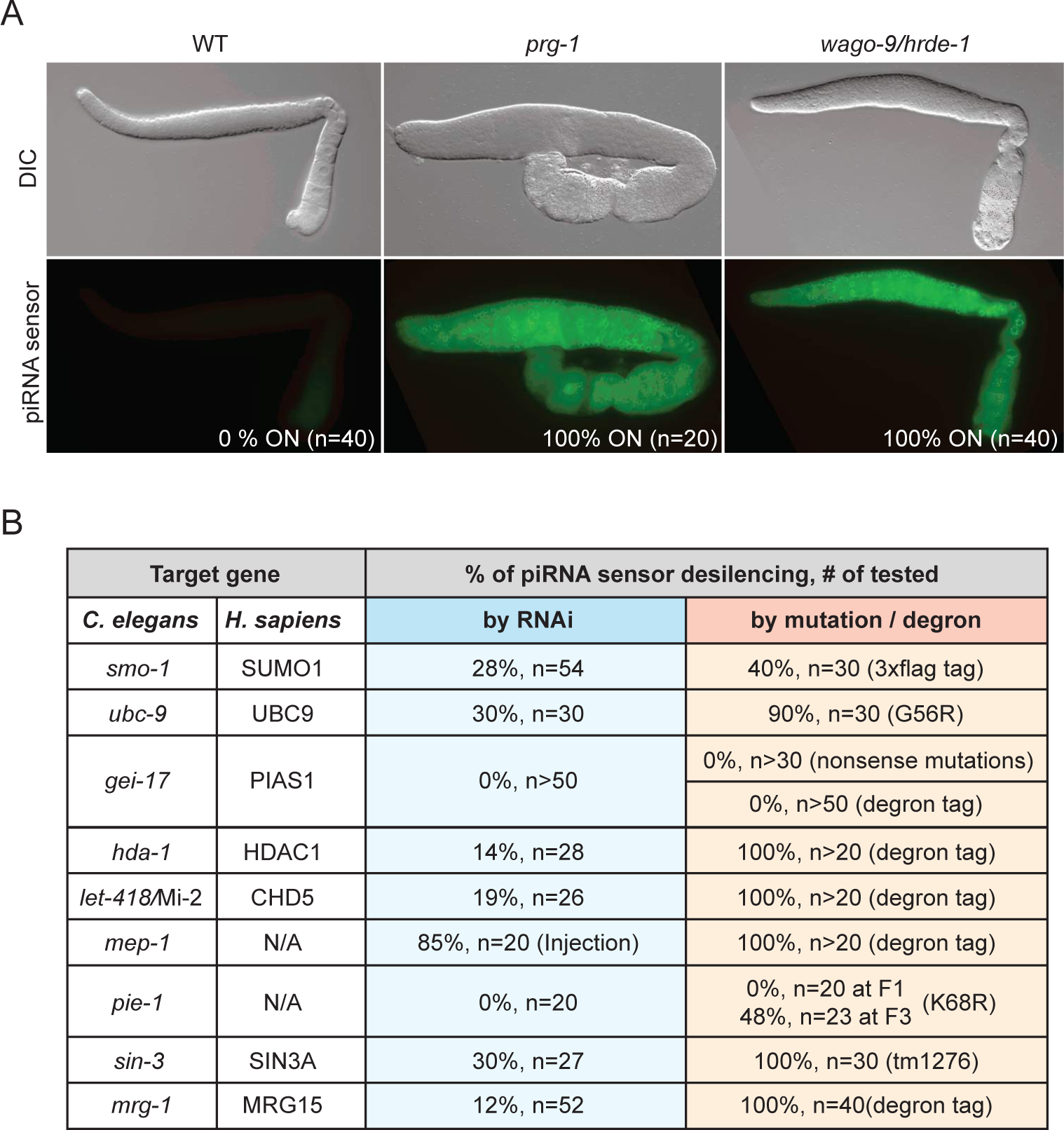
The SUMO pathway and chromatin remodeling factors promote piRNA silencing. (A) DIC and epifluorescent images of the dissected gonads showing piRNA sensor is desilenced in *prg-1(tm872)* and *wago-9/hrde-1(ne4769)*. (B) piRNA silencing requires SUMO and chromatin remodeling pathway. The desilenced GFP::CSR-1 transgenes (the p-granule signal) were scored in RNAi, mutants, or degron-auxin inducible protein depletion of the indicated target genes.

We were intrigued to find that silencing depended on SUMO (SMO-1), the SUMO conjugating enzyme (UBC-9), and components of known HDAC complexes (Figure 1B and Table S1). For example, depletion of the krüppel-type zinc finger protein-encoding gene *mep-1* and other NuRD-complex co-factors (*let-418*/Mi-2, *hda-1/*HDAC1, *lin-40*/MTA2/3, *lin-53*/RBBP4/7, and *dcp-66*/GATAD2B) all desilenced the piRNA sensor (Figure 1B and Table S1). Depletion of *sin-3/Rpd3* and *mrg-1/MORF4L1*, components of the SIN3-HDAC complex, also desilenced the reporter (Figure 1B and Table S1; see Discussion). Surprisingly, we found that depletion of the E3 SUMO ligase GEI-17 (PIAS1/Su(var)2-10) did not appear cause desilencing (Figure 1B).

To confirm that SUMO and HDAC factors are required for silencing of the piRNA sensor we selected genes from each pathway for further genetic study. Null alleles of many of these genes cause embryonic arrest, which precludes an analysis of silencing in the adult germline. We therefore tested whether partial or conditional loss-of-function alleles activate the piRNA sensor. For example, we found that a *3xflag::smo-1* gene fusion, which behaves like a partial loss-of-function allele of *smo-1*, desilenced the sensor in ∼40% of worms (Figure 1B), and a strain engineered to express the temperature sensitive UBC-9(G56R) protein (Kim et al., parallel) desilenced 90% of worms at the semi-permissive temperature of 23°C (Figure 1B). Strikingly, auxin-inducible degron alleles of *hda-1/*HDAC1*, mep-1, let-418/*Mi-2, and *mrg-1/*MRG15, and a viable null allele of *sin-3(tm1276)*, all caused 100% desilencing of the piRNA sensor (Figure 1B). By contrast, CRISPR generated auxin-inducible degron, as well as presumptive null alleles of the Su(var)2-10 paralog, *gei-17,* failed to de-silence the sensor strain (Figure 1B).

In a parallel study, we showed that the essential germline factor PIE-1 interacts with SUMO and UBC-9 and that SUMOylation of PIE-1 in the adult germline promotes SUMOylation and the activation of the type-1 HDAC, HDA-1 (Kim et al., parallel). We therefore tested whether a *pie-1* SUMO-acceptor site mutant PIE-1(K68R) (Kim et al., parallel) causes desilencing of the piRNA sensor. This homozygous viable mutant, which expresses a partially functional protein, initially silenced the piRNA sensor, but by the F3 generation, 48% of PIE-1(K68R) animals exhibited desilencing (Figure 1B), suggesting that PIE-1 SUMOylation promotes the trans-generational silencing of piRNA targets. Together, these findings suggest that histone deacetylase complexes are essential for piRNA-dependent silencing and that the SUMO pathway and PIE-1 (but not the E3 ligase homolog GEI-17) promote their function.

### SUMOylation of HDA-1 promotes piRNA surveillance

To test if the SUMO pathway promotes piRNA silencing via HDA-1 SUMOylation, we mutated SUMO-acceptor lysine residues on HDA-1. Human HDAC1 is SUMOylated near its C-terminus on lysines 444 and 476 (Figure 2A)(David et al., 2002). Despite poor conservation between the C-termini of human HDAC1 and worm HDA-1, we identified lysines 444 and 459 of HDA-1 as possible SUMO-acceptor sites (Figure 2A). To investigate whether one or both of these lysine residues is SUMOylated, we used CRISPR genome editing to mutate the endogenous *hda-1* gene in a strain that expresses SUMO fused to ten N-terminal histidines, *10xhis::smo-1* (Kim et al., parallel). We then captured SUMOylated proteins from worm lysates by nickel-nitrilotriacetic acid (Ni-NTA) affinity chromatography (Tatham et al., 2009) and analyzed eluates by western blotting for HDA-1. SUMOylated HDA-1 was robustly recovered from wild-type lysate and from lysates of each single-site lysine-to-arginine mutant, but was absent when both K444 and K459 were mutated together, HDA-1(KKRR) (Figure 2B). The protein MRG-1, which is highly SUMOylated in wildtype worms (Kim et al., parallel) remained highly SUMOylated in animals expressing HDA-1(KKRR) (Figure 2B).

**Figure 2.**
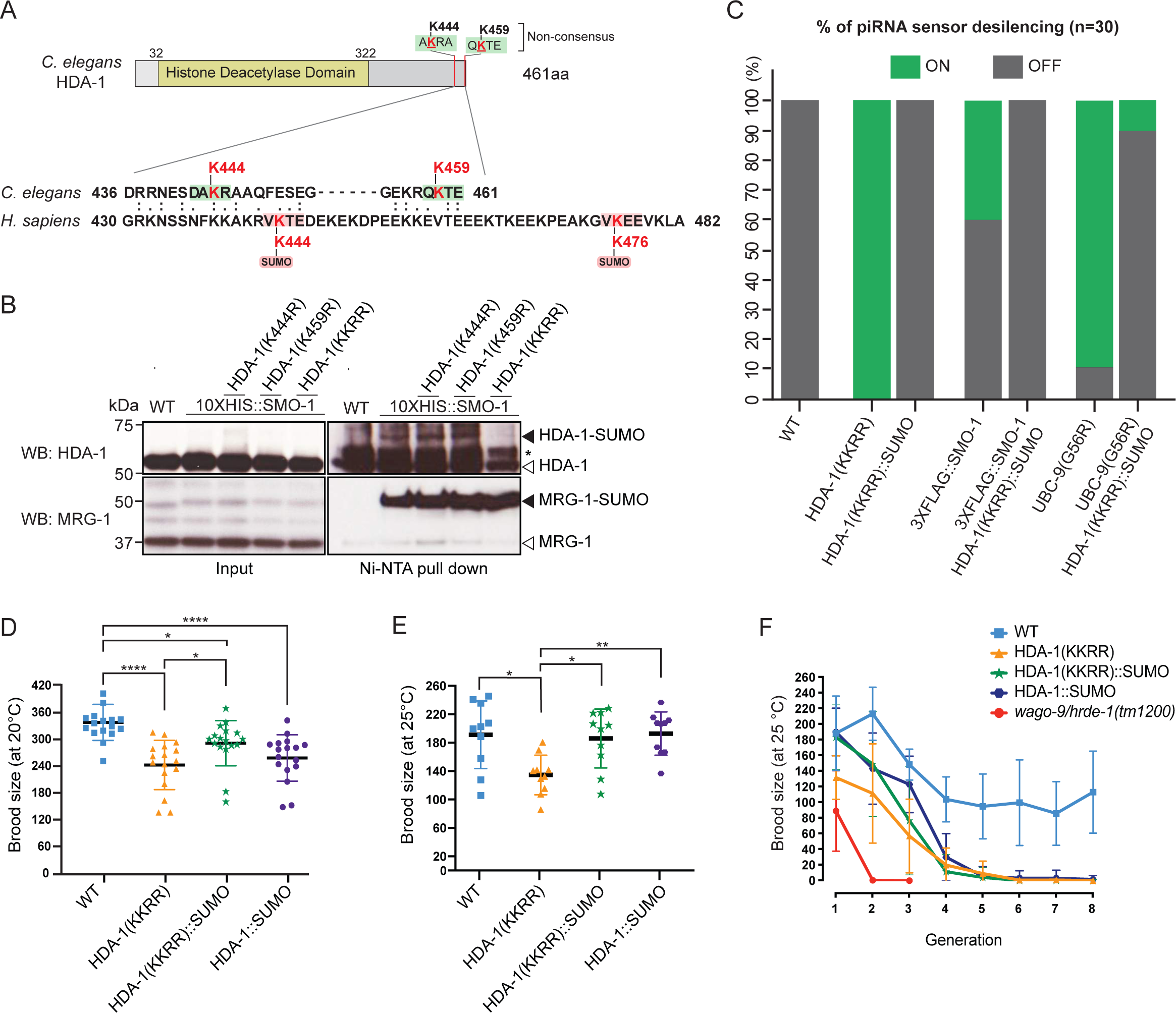
HDA-1 is SUMOylated on two residues K444 and K459 in *C. elegans*, which facilitates piRNA silencing. (A) Schematic representation of the SUMO acceptor sites in the *C. elegans* HDA-1 protein and partial sequence alignments between human HDAC1 and *C. elegans* HDA-1. In human, K444 and K476 are the two major SUMOylation sites (Red) within the putative SUMO consensus motif (Pink box). In *C. elegans*, two candidate SUMOylation sites, K444 and K459 (Red), but not consensus motifs (green box), are located on the C-terminus as indicated. (B) HDA-1 is SUMOylated on both K444 and K459 residues. *In vivo* SUMO purification followed by western blotting in adult lysates. The solid arrowhead indicates SUMOylated HDA-1 and MRG-1. Unmodified forms of HDA-1 and MRG-1 are indicated by the empty arrowhead and the asterisk (*) indicates a non-specific band for HDA-1 antibody. (C) HDA-1 SUMOylation is required for piRNA silencing. The desilenced piRNA sensor (expression of GFP::CSR-1 transgenes in p-granule) were scored in the wild-type (WT), HDA-1(KKRR), HDA-1(KKRR)::SUMO, 3xFLAG::SMO-1, UBC-9(G56R), 3xFLAG::SMO-1; HDA-1(KKRR)::SUMO, UBC-9(G56R); HDA-1(KKRR)::SUMO strains as indicated (%, n=30). (D and E) Brood sizes of WT, HDA-1(KKRR), HDA-1(KKRR)::SUMO, and HDA-1::SUMO at 20°C and at 25°C. Ordinary one-way ANOVA: *P<0.05, **P<0.01, ****<0.0001. (F) For the germline mortal assay, brood sizes of indicated strains (WT, HDA-1(KKRR), HDA-1(KKRR)::SUMO, HDA-1::SUMO, and *wago-9/hrde-1(tm1200)*) were scored over generations *at 25* °C. n=10, +/-s.e.m.

To determine if the HDA-1 lysine mutations affect piRNA-dependent silencing, we introduced the piRNA sensor into the HDA-1 SUMO-acceptor site mutants. Whereas the piRNA sensor remained silent in the *hda-1* SUMO single-site mutants (n=30), it was activated in 100% of HDA-1(KKRR) worms (Figure 2C). Strikingly, the silencing defect was completely rescued by inserting a modified *smo-1* coding sequence immediately before the stop codon of the *hda-1* SUMO-acceptor mutant, resulting in an HDA-1(KKRR)::SUMO translational fusion (Figure 2C; see Materials and Methods). Moreover, expression of HDA-1(KKRR)::SUMO rescued piRNA-dependent silencing in *smo-1* and *ubc-9* mutants (Figure 2C). These results strongly suggest that the SUMO pathway promotes piRNA silencing via C-terminal SUMOylation of HDA-1.

### HDA-1 SUMOylation mutants cause temperature-dependent reductions in fertility

Factors that promote genome integrity and epigenetic inheritance are required for germline immortality: their loss causes a mortal germline phenotype, whereby fertility declines in each generation and this decline is often exacerbated at elevated temperatures (Ahmed and Hodgkin, 2000). Although *hda-1* is an essential gene required for embryonic development (Shi and Mello, 1998), worms expressing HDA-1(KKRR) and HDA-1(KKRR)::SUMO from the endogenous *hda-1* locus were viable and fertile. Careful examination of brood size revealed that HDA-1(KKRR) animals made significantly fewer progeny than wild-type worms at 20°C and at 25°C (Figures 2D and 2E). Appending SUMO via translational fusion, HDA-1(KKRR)::SUMO rescued this fertility defect, suggesting that HDA-1 SUMOylation promotes fertility (Figures 2D and 2E). The fertility of worms expressing HDA-1(KKRR) steadily declined over several generations when animals were maintained at 25°C, from an average of 132 progeny in the first generation to fewer than ten progeny in the fifth and subsequent generations (Figure 2F). Wild-type worms also showed an initial decline in fertility when maintained at 25°C, but averaged ∼100 progeny in the 4^th^ and subsequent generations. By contrast, *wago-9/hrde-1* mutants showed a rapid decline in fertility and could not be maintained beyond the 3^rd^ generation (Figure 2F) (Buckley et al., 2012; Spracklin et al., 2017). Interestingly, although HDA-1(KKRR)::SUMO animals exhibited wild-type fertility in the first generation at 25°C, fertility gradually declined over five generations at 25°C (Figure 2F). Worms expressing wild-type HDA-1 fused to SUMO, HDA-1::SUMO showed a similar generational loss in fertility (Figure 2F). Thus the regulated SUMOylation of HDA-1 is essential for germline immortality.

### SUMOylation promotes HDA-1 association with other chromatin factors including NuRD complex components

SUMOylation modulates protein interactions (Hendriks and Vertegaal, 2016; Kerscher, 2007). We therefore wished to examine how SUMOylation affects HDA-1 complexes. We introduced a GFP tag into the C-terminus of the endogenous wild-type and mutant *hda-1* alleles, and used GFP-binding protein (GBP) beads to immunoprecipitate the HDA-1::GFP fusion proteins (Rothbauer et al., 2008). SDS-PAGE analysis revealed that a core set of proteins strongly interact with HDA-1::GFP (Figure 3A). Mass-spectrometry (MS) of the corresponding gel slices identified these proteins as: LIN-40, a homolog of metastasis-associated protein (MTA1) (Chen and Han, 2001); LIN-53, a homolog of retinoblastoma-associated protein 46/48 (RBAP46/48) (Solari and Ahringer, 2000); DCP-66, a homolog of GATA zinc-finger domain containing protein GATAD (Kaser-Pebernard et al., 2014); and SPR-5, a homolog of lysine demethylase (LSD1/KDM1) (Katz et al., 2009).

**Figure 3.**
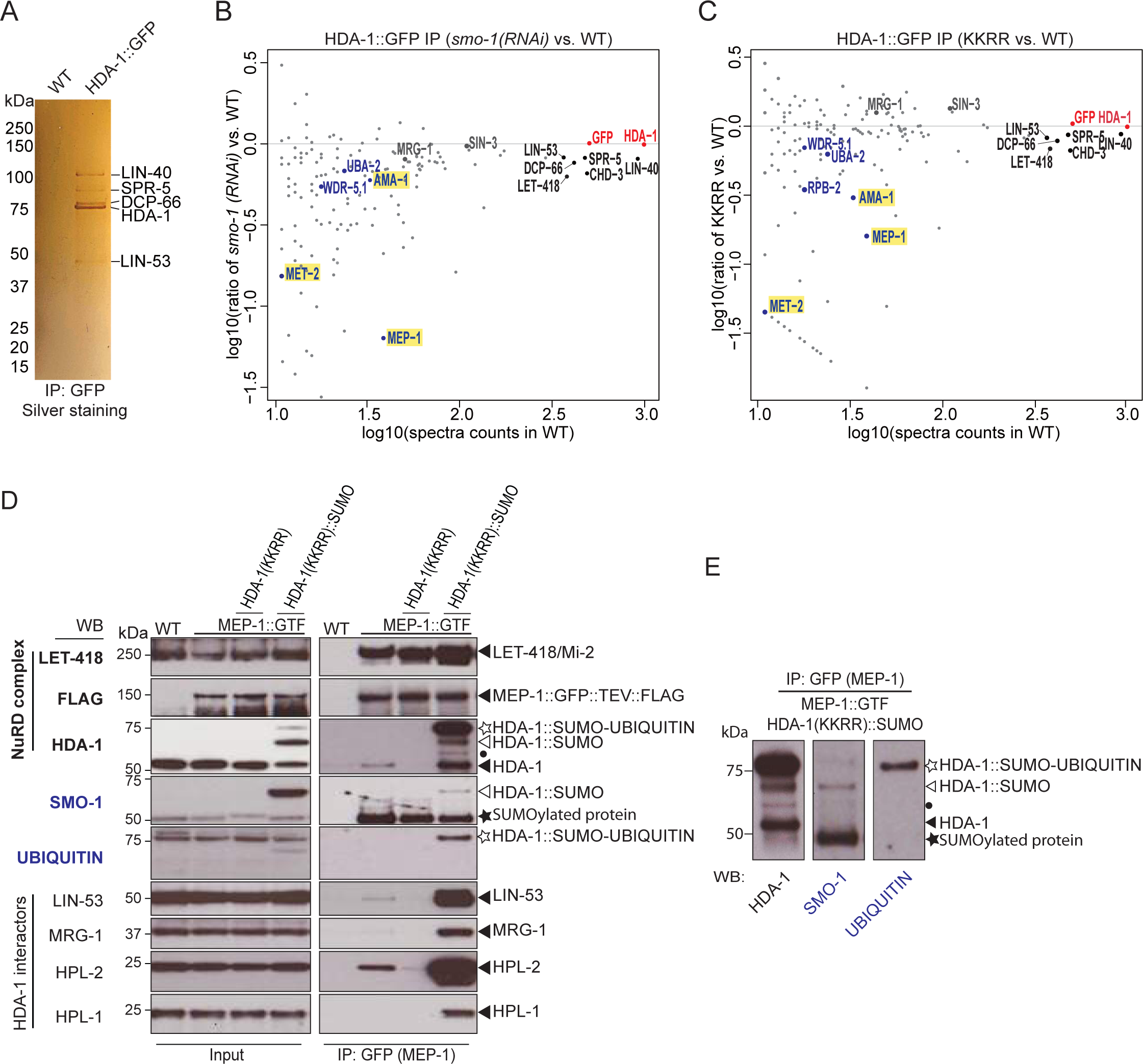
SUMO pathway promotes HDA-1 association with other chromatin factors including NuRD complex component, MEP-1. (A) Silver staining of GFP IP in WT and *hda-1::gfp*. The Five apparent protein bands were cut and analyzed by Mass spectrometry separately and the major protein identified in each band was labeled aside. (B and C) Scatter plot of HDA-1 GFP IP-MS results. X axis is the log value of protein spectra counts in WT worms; Y axis is the log ratio of spectra counts (B) in *smo-1* RNAi treated worms versus WT worms and (C) in mutants expressing HDA-1(KKRR) versus WT worms. The spectra count of HDA-1 was used to normalize between samples. For the space reason, some proteins of interest were labeled. Please see Table S2 for all proteins identified. (D and E) HDA-1 SUMOylation promotes the associate of MEP-1 with chromatin regulators. MEP-1 GFP IP followed by western blot against the indicated antibody. The black arrowheads indicate the specific bands for the corresponding antibodies. The white arrowheads, white star, and black dot/star indicate the HDA-1(KKRR)::SUMO, HDA-1(KKRR)::SUMO-Ubiquitin, and unknown HAD-1 isoform/unknown SUMOylated protein band, respectively. In (E), the three lanes were from the same membrane and well aligned based on protein size. Protein names of these bands were labeled.

We also used Reverse phase high-performance liquid chromatography, RP-HPLC, mass spectrometry to identify HDA-1::GFP interactors (see Materials and Methods). Among approximately 200 very high confidence interactors with ≥10 spectral counts, we identified 63 that were depleted by an arbitrary cut off of 40% in immunoprecipitates from both *smo-1(RNAi)* and HDA-1(KKRR)::GFP lysates (Figures 3B and 3C and Table S2).

These SUMO-dependent interactors included MEP-1 (Unhavaithaya et al., 2002), AMA-1 (major subunit of pol lI) (Sanford et al., 1983), and MET-2 (SETDB1 H3K9 histone methyltransferase) (Andersen and Horvitz, 2007; Bessler et al., 2010). This analysis also revealed that the core interactors identified above, LIN-40, LIN-53, DCP-66, and SPR-5, were also reduced (by 12% to 37%) in IPs from *smo-1(RNAi)* or HDA-1(KKRR) lysates (Figures 3B and 3C and Table S2). Of note, interactions between HDA-1 and SIN-3/SIN3A were not sensitive to SUMO-pathway perturbations (Figures 3B and 3C and Table S2).

### HDA-1 SUMOylation promotes the association of MEP-1 with chromatin regulators

To explore the molecular consequences of HDA-1 SUMOylation on its physical interactions with MEP-1 and other chromatin regulators, we crossed a tandemly tagged *mep-1::gfp::tev::3xflag* allele (MEP-1::GTF; Kim et al., parallel) into animals expressing HDA-1(KKRR) and HDA-1(KKRR)::SUMO proteins and then used GBP beads to purify MEP-1::GTF protein complexes from otherwise wild-type or *hda-1* mutant lysates. As expected, interactions between MEP-1::GTF and LET-418/Mi-2 did not depend on HDA-1 SUMOylation (Figure 3D and Kim et al., parallel). Consistent with our HDA-1 proteomics studies, however, MEP-1::GTF pulled down wild-type HDA-1 but not HDA-1(KKRR) (Figure 3D). Moreover, LIN-53/RBBP4/7, MRG-1/MRG15, and HPL-1/-2/HP1 were also greatly reduced in MEP-1 complexes purified from HDA-1(KKRR) lysates (Figure 3D). Strikingly, each of these factors were dramatically increased, above wild-type levels, in MEP-1 complexes purified from HDA-1(KKRR)::SUMO lysates (Figure 3D). Thus, the C-terminal fusion of SUMO to HDA-1 rescues MEP-1 interactions with HDA-1(KKRR), and also promotes MEP-1 interaction with HDA-1-binding partners.

In MEP-1 immunoprecipitates from HDA-1(KKRR)::SUMO lysates, we detected multiple HDA-1 bands, including a prominent band slightly larger than the expected size of HDA-1::SUMO (white stars in Figures 3D and 3E). Western blots with antibody specific to Ubiquitin suggested that this prominent band is a mono-ubiquitinated form of the fusion protein (Figures 3D and 3E). In both input and IP samples, we also observed an isoform similar in size to endogenous HDA-1 that was not detected by SUMO-specific antibodies in our western blot assays (Figures 3D and 3E), suggesting that the HDA-1::SUMO fusion protein may be cleaved near the C-terminus of HDA-1, removing the SUMO peptide.

### HDA-1 SUMOylation promotes histone deacetylation *in vivo*

The findings above suggest that SUMOylation promotes the assembly and function of HDA-1 complexes. Our proteomics studies also revealed that SUMOylation promotes interactions between HDA-1 and other histone modifying enzymes required for heterochromatin formation, including the demethylase SPR-5/LSD1, which removes activating H3K4me2/3 marks, and the methyltransferase MET-2/SetDB1, which installs silencing H3K9me2/3 marks (Greer and Shi, 2012). Consistent with the idea that these factors promote silencing, immunostaining of adult hermaphrodite gonads from HDA-1(KKRR) animals and from *mep-1*-depleted worms revealed dramatically higher levels of H3K9Ac compared to wild-type (Figure 4A and Figure S1A). In addition, increased levels of active chromatin H3K4me3 and much lower levels of the silencing mark H3K9me2 were detected throughout the germline compared to wild-type (Figures S1B-S1C and Figures 4C-4D). Moreover, we found that HDA-1, LET-418/Mi-2, and MEP-1 (including MEP-1 expressed from the germline-specific *wago-1* promoter) bind heterochromatic regions of the genome, depleted of the activating H3K9Ac mark and enriched for the silencing marks H3K9me2/3 (Figure 4B). Thus, SUMOylation of HDA-1 appears to drive formation or maintenance of germline heterochromatin.

**Figure 4.**
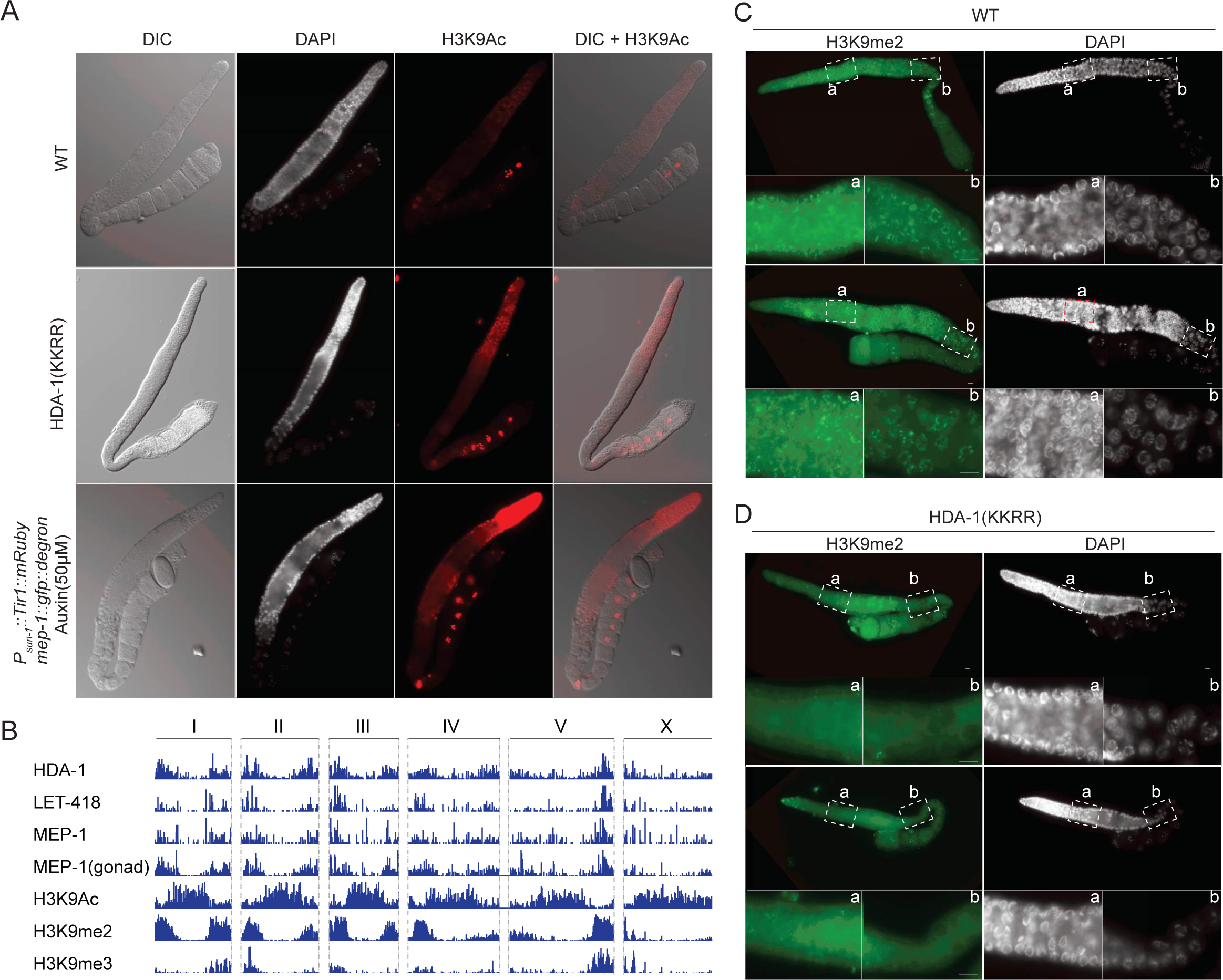
HDA-1 SUMOylation is required for its histone deacetylase activity and silenced chromatin state. (A) DIC and immunofluorescence micrographs of anti-H3K9Ac and DAPI staining in adult gonad of WT, HDA-1(KKRR) mutants, *mep-1::gfp::degron* with auxin (50*μm*). (B) IGV view of ChIP-seq peaks of NuRD complex members (HDA-1, LET-418, and MEP-1) and three histone marks (H3K9Ac, H3K9me2, and H3K9me3) generated by MACS2. The whole genome of *C. elegans* is shown and the Chromosome number is labeled at the top. MEP-1(gonad) stands for the ChIP experiments from a germline specifically expressed *mep-1::gfp::3xflag* driven by a *wago-1* promoter. (C and D) Immunofluorescence micrographs of anti-H3K9me2 and DAPI staining in adult gonad of (C) WT and (D) HDA-1(KKRR). The dashed boxes indicated as “a” and “b” were enlarged as shown.

### SUMOylated HDA-1 and PRG-1 co-regulate hundreds of targets, including many spermatogenesis genes

The visibly reduced levels of germline heterochromatin caused by the loss of HDA-1 SUMOylation suggests that gene expression is broadly misregulated when HDA-1 function is compromised. To further examine the impact of the HDA-1 and SUMO pathways on germline gene expression, we performed high-throughput sequencing of mRNAs isolated from dissected germlines of HDA-1(KKRR), HDA-1(KKRR)::SUMO, and UBC-9(G56R) animals, and from degron alleles of *hda-1* and *mep-1* with or without auxin exposure beginning at the L4 stage (Figure 5A). For each mutant, replicate libraries gave highly reproducible mRNA profiles (Figures S2C-S2M). Depletion of HDA-1 or MEP-1 or inactivation of the SUMO pathway caused a wide-spread upregulation of germline mRNAs and transposon families, with extensive but incomplete overlap between the mutants (Figure 5B and Figures S2A, S3A, and S3B). Twice as many genes were upregulated in *degron::hda-1* germline as in *degron::mep-1* or UBC-9(G56R) (Figure 5B and Figure S2A), likely reflecting the role of HDA-1 in multiple complexes. Nearly 10-fold fewer genes were upregulated in HDA-1(KKRR) than in auxin-treated *degron::hda-1* animals (Figure 5B and Figure S2A), consistent with the phenotypic differences between the two mutants. Most (305, ∼71%) of the genes upregulated in HDA-1(KKRR) germlines were also upregulated auxin-treated *degron::hda-1* animals (Figure 5B). As expected, mRNAs upregulated in HDA-1(KKRR) were restored to nearly wild-type levels in HDA-1(KKRR)::SUMO (Figures 5C and 5D). Consistent with the known role of the NuRD complex and SUMOylation pathways in modulating chromatin states, we observed a loss of enrichment for H3K9me2, as measured by ChIP-seq, near the promoters of genes upregulated in HDA-1(KKRR), UBC-9(G56R), and auxin-treated *degron::hda-1* worms (Figures S4A-S4D).

**Figure 5.**
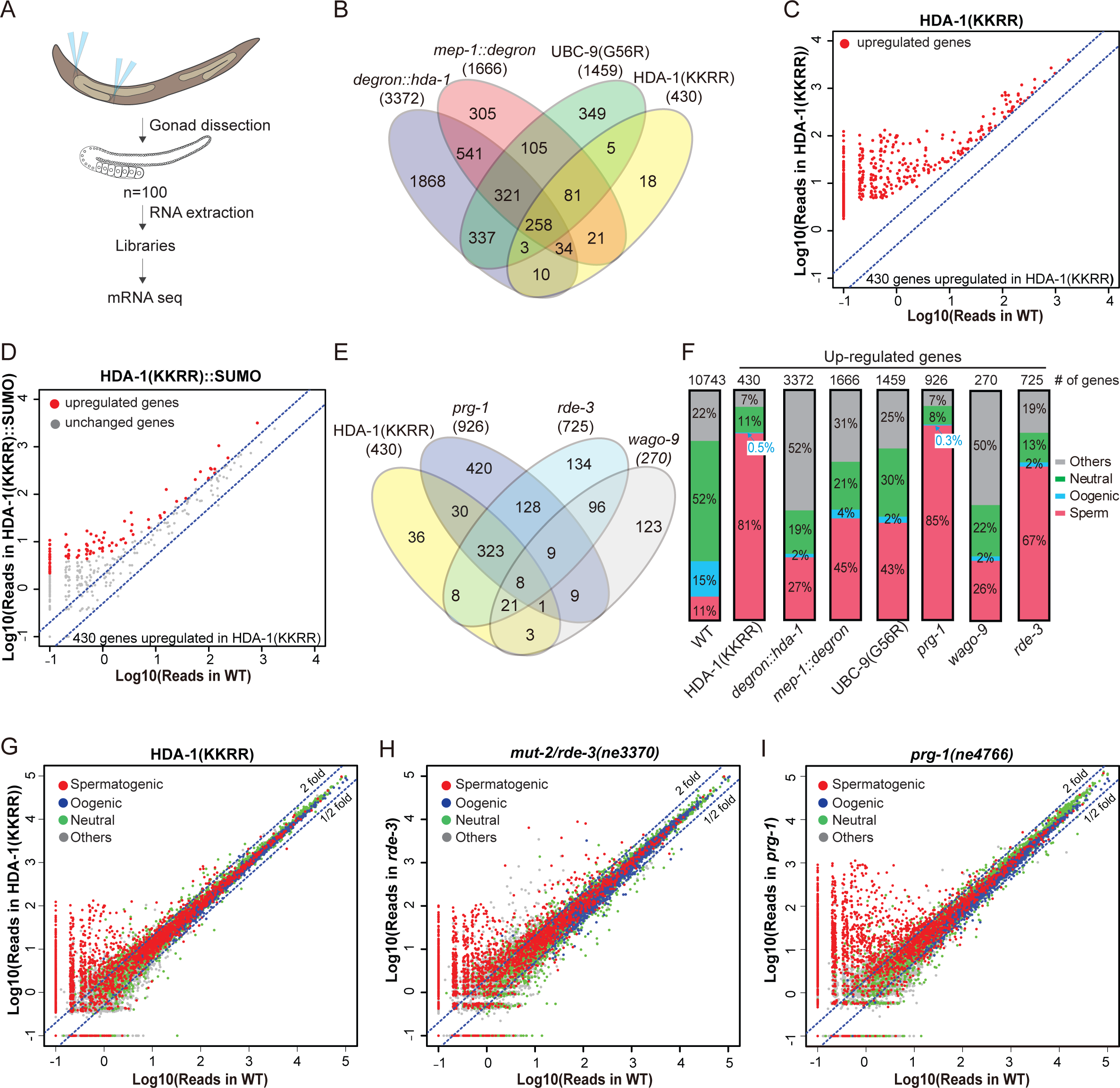
SUMO pathway, the NuRD complex and piRNA pathway are regulating the same group of targets through HDA-1 SUMOylation. (A) Schematic showing mRNA-seq from the dissected gonads. (B) Venn diagram shows the overlap of upregulated genes in *degron::hda-1*, *mep-1::degron*, UBC-9(G56R) and HDA-1(KKRR). The total numbers of the upregulated genes in each mutant are indicated in the parentheses. (C) Scatter plot of genes upregulated in HDA-1(KKRR). X axis is the reads in WT, and Y axis is the reads in HDA-1(KKRR). (D) Scatter plot of genes upregulated/unchaged in HDA-1(KKRR)::SUMO among 430 upregulated genes in HDA-1(KKRR) in (C). X axis is the reads in WT, and Y axis is the reads in HDA-1(KKRR)::SUMO. (E) Venn diagram showing the overlap of upregulated genes between HDA-1(KKRR), *prg-1, rde-3* and *wago-9* mutants. (F) Bar graph shows the composition of each category in each mutant. “Others” are the ones cannot be put into these categories based on the previous report (Ortiz et al., 2014). Genes with read count more than 1 in the mRNA seq of WT gonad were used to generate the “WT gonad” dataset as a reference (total gene number is 10743). The number of upregulated genes in each mutant is labeled at the top. (G and I) Scatter plot of reads (G) in HDA-1(KKRR), (H) *rde-3*, and (I) *prg-1(ne4766)* versus those in WT. Red, blue and green dots are the genes annotated as “Sperm” (shorten “Spermatogenic”), “Oogenic” and “Neutral”, respectively. The blue dashed lines indicate two-fold increase or decrease versus WT. A value of 0.1 was assigned to the missing value, thus the ones with a X value of “-1” were not detected in WT.

Because HDA-1 SUMOylation is required for silencing of a piRNA reporter, we wished to explore what endogenous mRNAs are co-regulated by HDA-1 SUMOylation and the piRNA pathway factors, PRG-1, WAGO-9/HRDE-1, and RDE-3. To address this question we prepared mRNA sequencing libraries from dissected gonads collected from *prg-1(ne4766), wago-9/hrde-1(tm1200)*, and *rde-3(ne3370)* mutant animals (Figure 5A). RDE-3 is required for RdRP-dependent amplification of small RNAs (termed 22G-RNAs) that guide transcriptional and post-transcriptional silencing by WAGO Argonautes (Gu et al., 2009; Zhang et al., 2011). This RDE-3-dependent WAGO-RdRP system can function independently of upstream Argonautes once initiated and plays a role in the amplification and maintenance of silencing triggered by both the piRNA and dsRNA-induced silencing pathways (Shirayama et al., 2012). Of the 430 mRNAs upregulated (≥2-fold) in HDA-1(KKRR), we found that 362 (84%) were upregulated (≥2-fold) in *prg-1(ne4766)*, 360 (84%) were upregulated in *rde-3(ne3370)*, and 331 (77%) were upregulated in all three mutant strains (Figure 5E). Similarly, the genes upregulated in HDA-1(KKRR) accounted for 40% of the genes upregulated in *prg-1(ne4766)* and 50% of those in *rde-3(ne3370)*

(Figure 5E). Fewer genes were upregulated in *wago-9/hrde-1* (Figure 5E and Figure S2A), perhaps due to redundancy with other WAGO Argonautes. Whereas only five transposon families were upregulated in *prg-1(ne4766)* (Figures S3A and S3B), thirty were upregulated in *rde-3* mutants (Figures S3A and S3B). These latter findings are consistent with the observation that once initiated, piRNA-induced silencing can be maintained by the WAGO Argonaute-RdRP system and chromatin silencing machinery, independently of piRNAs or PRG-1 (Shirayama et al., 2012). A total of six transposon families were upregulated in HDA-1(KKRR) (Figures S3A and S3B). Comparison with mRNA sequencing data from auxin-treated *degron::hda-1* animals revealed an even more extensive overlap with Piwi pathway mutants (Figure S2B), indicating that HDA-1 also promotes target silencing independently of HDA-1 SUMOylation.

Most of the genes upregulated in *prg-1*, *rde-3*, and HDA-1(KKRR) mutants are normally expressed during spermatogenesis (Figures 5F-I and Figures S5A and S5B). In most cases, the upregulation of these spermatogenesis mRNAs did not correlate with reduced RdRP-derived 22G-RNAs targeting these genes (Figures S6A and S6B), suggesting that the spermatogenesis switch may be regulated independently of (or indirectly by) the WAGO 22G-RNA pathway (See Discussion).

### HDA-1 physically interacts with WAGO-9/HRDE-1 and functions in inherited RNAi

Because WAGO-9/HRDE-1 is a nuclear Argonaute that functions downstream in the Piwi pathway to maintain epigenetic silencing (Ashe et al., 2012; Bagijn et al., 2012; Buckley et al., 2012; Shirayama et al., 2012), we asked if WAGO-9/HRDE-1 interacts with HDA-1. We used CRISPR genome editing to generate a functional *gfp::wago-9* strain and then used GBP beads to immunoprecipitate GFP::WAGO-9 complexes. Western blot analyses revealed that HDA-1 co-precipitates specifically with GFP::WAGO-9 (Figure 6A). We also found that the HP1-like protein HPL-2 interacts with WAGO-9/HRDE-1 (Figure 6A), consistent with previous genetic studies (Ashe et al., 2012; Buckley et al., 2012; Gu et al., 2012; Luteijn et al., 2012; Shirayama et al., 2012). HPL-2 binds methylated forms of H3K9 in heterochromatin (Garrigues et al., 2015). Neither HDA-1 nor HPL-2 were precipitated by GBP beads incubated with lysates prepared from untagged wild-type worms. These interactions were confirmed in reciprocal GFP-IP experiments on *hda-1::gfp* lysates (Figure 6B). Moreover, in these latter studies, we found that the interaction between WAGO-9 and HPL-2 was reduced in HDA-1(KKRR) animals or by *smo-1(RNAi)* (Figure 6B). These findings suggest SUMOylation of HDA-1 promotes the assembly of an Argonaute-guided nucleosome remodeling complex.

**Figure 6.**
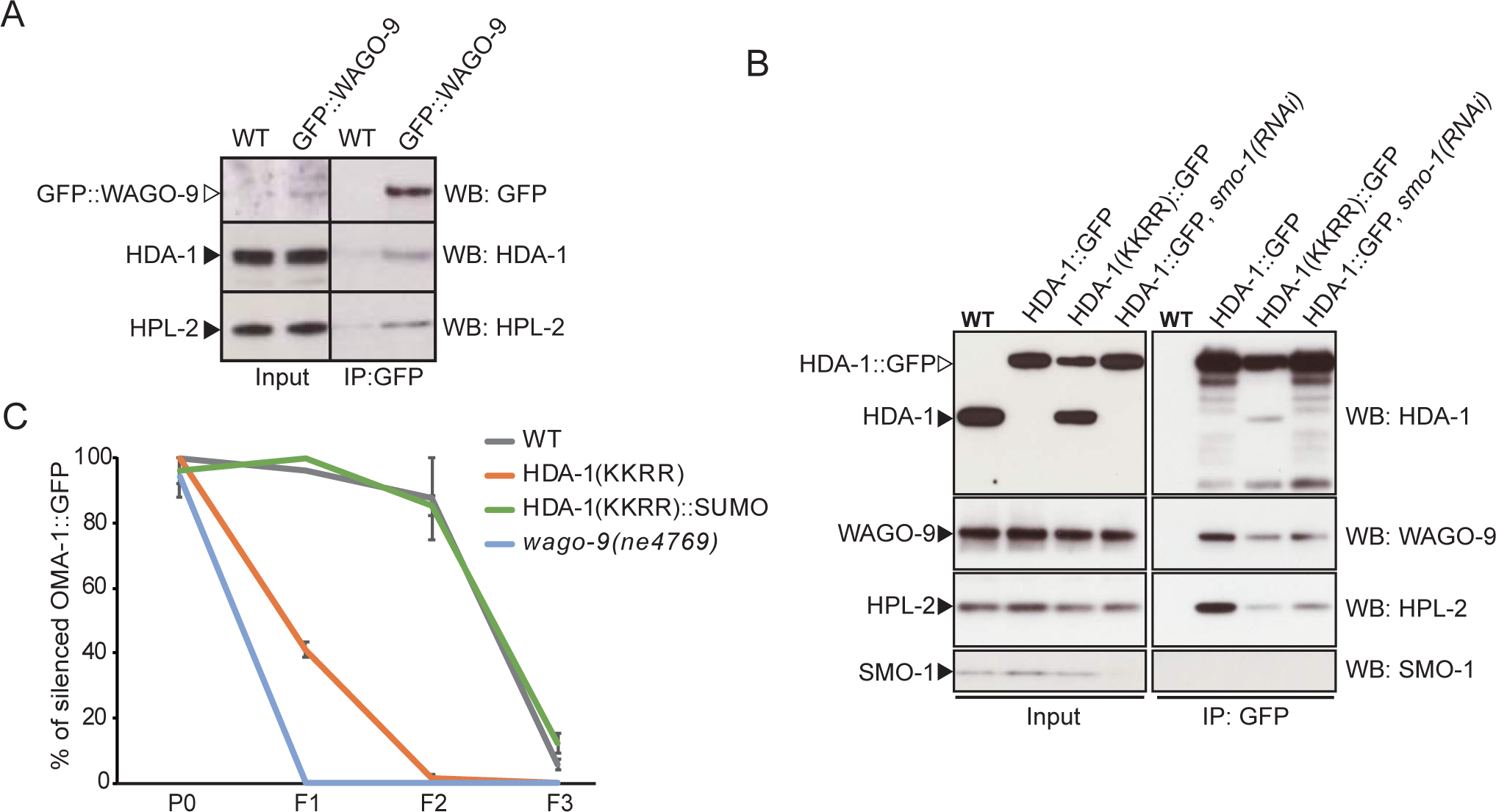
HDA-1 interacts with WAGO-9/HRDE-1, which depends on HDA-1 SUMOylation. (A) GFP::WAGO-9/HRDE-1 IP followed by western blotting. GFP IP was done with synchronized young adult worms and IP samples were blotted with GFP, HDA-1 and HPL-2 antibodies. (B) HDA-1::GFP IP followed by western blotting. HDA-1 GFP IP samples were blotted with HDA-1, WAGO-9/HRDE-1, HPL-2 antibodies. Blotting with anti-SMO-1 showed depletion of SMO-1 in the *smo-1* RNAi treated samples. The band of the same size as endogenous HDA-1 (solid arrowhead) in HDA-1(KKRR)::GFP sample probably indicates the cleavage product and the cleavage site is close to the C-terminus of HAD-1. (C) HDA-1 SUMOylation functions in RNAi inheritance. The indicated strains were treated with *gfp* RNAi at P0 and transferred to regular NGM plates from F1. The percentage of worms with OMA-1::GFP was scored (N>=22 for two replicates). Standard error of the mean (SEM) was used for the error bar.

Many of the factors that maintain heritable epigenetic silencing triggered by piRNAs—including WAGO-9/HRDE-1—are also required for multigenerational germline silencing triggered by exogenous dsRNA, i.e., inherited RNAi (Ashe et al., 2012; Buckley et al., 2012; Shirayama et al., 2012). We therefore asked if HDA-1 SUMOylation is also required for inherited RNAi. Worms expressing a bright germline OMA-1::GFP were fed bacteria that express GFP dsRNA (i.e., RNAi by feeding) for one generation (P0), and subsequent F1 and F2 generations were fed the normal *E. coli* diet (no *gfp* dsRNA). In wild-type worms, *oma-1::gfp* was silenced in the P0 generation (in the presence of *gfp* dsRNA), and remained silent after the removal of *gfp* dsRNA in the F1 and F2 generations (Figure 6C). By contrast, as previously shown (Ashe et al., 2012; Buckley et al., 2012; Spracklin et al., 2017), although they efficiently and fully established RNA interference in the presence of dsRNA (the P0 generation), *wago-9/hrde-1* animals exhibited a full recovery of GFP expression in the F1 and F2 generations after the removal of dsRNA. We found that HDA-1(KKRR) animals were similarly defective for heritable RNAi: *oma-1::gfp* silencing was efficiently established in the P0 but upon dsRNA removal GFP expression was fully restored in the F1 and F2 generations (Figure 6C). Finally, we found that restoring SUMO to HDA-1 by translational fusion, in animals expressing HDA-1(KKRR)::SUMO completely restored heritable RNAi (Figure 6C). Taken together, these findings suggest that HDA-1 SUMOylation promotes heritable transcriptional silencing in both the piRNA and RNAi pathways.

## Discussion

Argonaute small RNA pathways collaborate with chromatin factors to co-regulate dispersed genetic elements and chromosomal regions (Reviewed in Almeida et al., 2019). This system, which likely evolved in part to hold in check the activity of transposons, has the added effect of wiring together and coordinating genome-wide networks of gene regulation. In doing so, it has been proposed that these mechanisms may also underlie the ability of multicellular organisms to efficiently and stably task their genomes to the diverse programs of gene expression that support multicellularity (Cowley and Oakey, 2013; Dubin et al., 2018; Ninova et al., 2020; Sundaram et al., 2014). Here we have used *C. elegans* to explore how Argonaute systems orchestrate the transition of an actively expressed gene to a stably silenced state. We have shown that the initiation of transcriptional silencing requires SUMOylation of conserved C-terminal lysine residues in the type-1 histone deacetylase HDA-1. The SUMOylation of HDA-1 promotes its activity, while also promoting physical interactions with other components of a germline nucleosome-remodeling histone deacetylase (NuRD) complex, as well as the nuclear Argonaute HRDE-1/WAGO-9 and the heterochromatin protein HPL-2 (HP1). Our findings suggest how the SUMOylation of HDAC promotes the co-assembly of a nuclear Argonaute complex that establishes repressive chromatin state for transcriptional silencing in response to dsRNA and piRNA cues (Figure 7).

**Figure 7.**
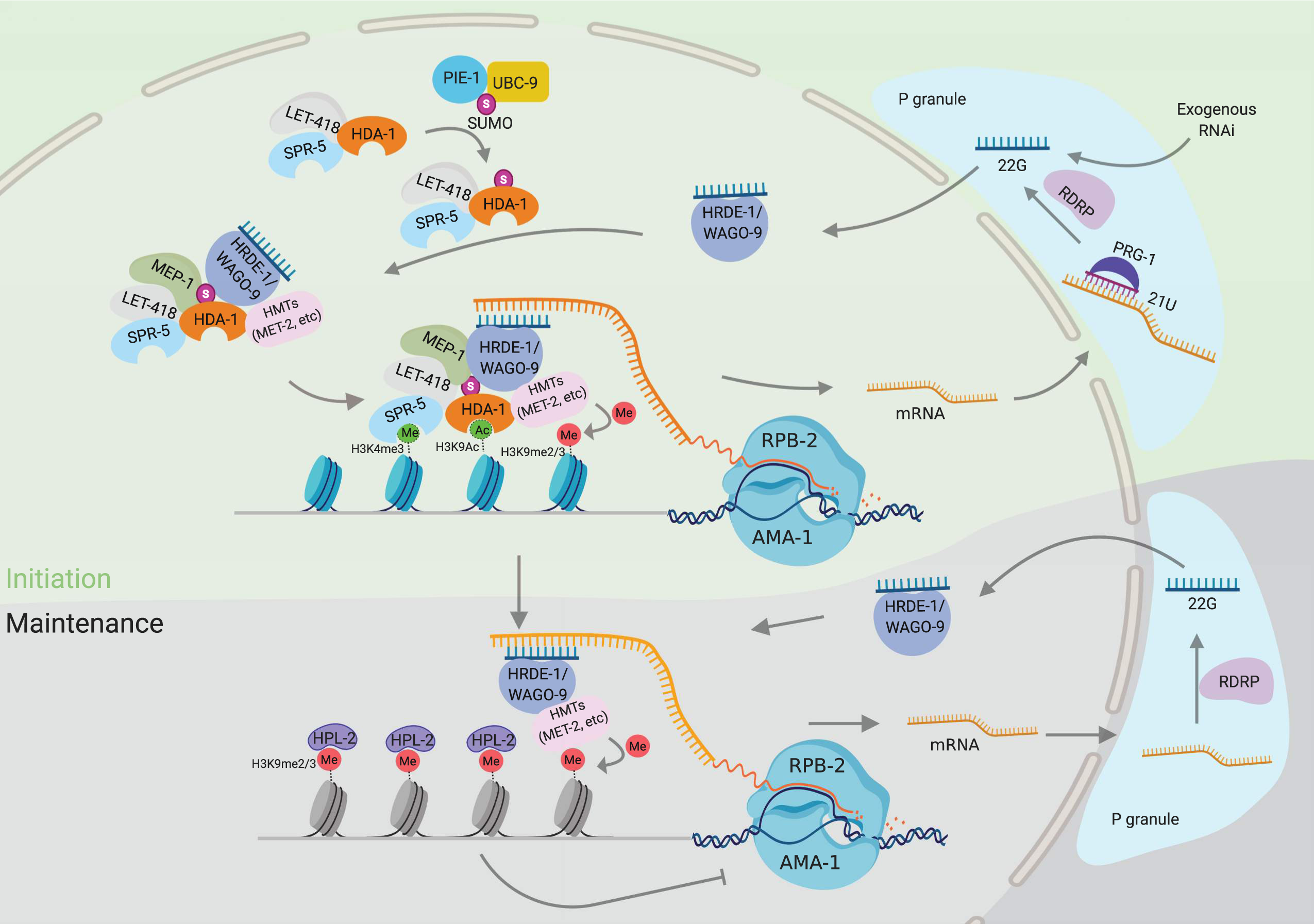
Model: HDAC1 SUMOylation promotes Argonaute directed transcriptional gene silencing. SUMOylation of HDA-1 enables the assembly of the NuRD complex to regulate its activity and helps the NuRD complex to be recruited to piRNA targets through WAGO-9 or other Argonautes. Together with other enzymes, like SPR-5, MET-2 *etc*., the huge histone remodeling machine removes the active histone marks (H3K9Ac and H3K4me2/3) and establishes the silence histone marks (H3K9me2/3) around the targets to suppress their transcription.

### Complementation of SUMO-acceptor lysine mutations by a SUMO translational fusion

SUMO, like other posttranslational modifications such as protein phosphorylation or ubiquitination, functions in numerous and diverse cellular mechanisms (Gareau and Lima, 2010; Psakhye and Jentsch, 2012; Rosonina et al., 2017). The multiple essential functions of the SUMOylation pathway make it nearly impossible to interpret the exact cause of phenotypes associated with mutations in SUMO or in other core components of the SUMOylation pathway. Even mutation of specific SUMO acceptor sites could cause confounding effects, potentially preventing other post-translational lysine modifications or disrupting protein folding or protein-binding interfaces. Moreover, mutation of a specific SUMO acceptor site often causes subtle or no phenotypic consequence (Psakhye and Jentsch, 2012; Sacher et al., 2006; Silver et al., 2011). This fact and the finding that many components of a complex are SUMOylated together has led to the idea that SUMO modifications can function synergistically, each contributing incrementally, for example, to stabilize a complex composed of multiple SUMOylated proteins that function together (Psakhye and Jentsch, 2012).

In this study, we employed a “complementation” strategy, first mutating the SUMO-acceptor lysines in HDA-1 and then inserting an in-frame SUMO coding sequence (lacking its C-terminal conjugation motif; Estruch et al., 2016) immediately upstream of the HDA-1 stop codon. We were surprised to find that the HDA-1::SUMO fusion was so well tolerated, despite dramatically increasing the association of MEP-1 and HDA-1. For example, HDA-1::SUMO did not interfere with the MEP-1-dependent silencing of germline gene expression in the embryonic soma, nor did it ectopically activate this germline-gene transcriptional silencing program in the adult germline. Instead, HDA-1::SUMO properly directed the stage-appropriate role of HDA-1 in Argonaute-dependent transcriptional silencing. The ability of the HDA-1::SUMO fusion to complement upstream mutations in the SUMO pathway, strongly indicates that the translationally fused SUMO peptide functions in a manner similar to the SUMO moieties that are normally conjugated to the C-terminal SUMO-acceptor lysines of HDA-1.

The *hda-1::sumo* fusion gene appeared to express several HDA-1 protein isoforms likely due to additional posttranslational modifications (Figures 3D and 3E). These included a mono-ubiquitinated isoform and a truncated isoform that appeared to lack SUMO altogether at its C-terminus. Interestingly, the mono-ubiquitinated fusion protein was dramatically enriched in the MEP-1 co-IP. We noticed that the ubiquitinated isoform exhibited dramatically reduced staining with a SUMO-specific antibody, raising the possibility that SUMO itself is modified by Ubiquitin as has been reported for SUMO isoforms in humans (Danielsen et al., 2011; Tatham et al., 2008). We have not yet explored the potential physiological significance of these findings. Perhaps the translational conjugation of SUMO is fairly well-tolerated in HDA-1 because the cleavage and/or ubiquitination of the fusion protein provide mechanisms to counter its activity. Such mechanisms could serve as additional safeguards that with SUMO-deconjugating enzymes allow reversal or modulation of SUMO-specific regulatory effects.

Several studies have investigated HDAC1 C-terminal lysine mutations (KKRR mutants) in mice and in human cell culture (Cheng et al., 2004; David et al., 2002; Gill, 2005); (Citro et al., 2013). While all of these studies revealed changes consistent with the likely importance of these residues in promoting HDAC1 functions, only one study explored whether appending SUMO to HDAC1 by translational fusion had opposing effects. Remarkably, this study showed that a lenti-virus driven of an HDAC1::SUMO fusion protein rescued the learning and memory deficits of an APP/Presenilin 1 murine model of Alzheimer’s disease (Tao et al., 2017). Our finding that conjugating SUMO to HDA-1(KKRR), by precision genome editing at the endogenous *hda-1* locus in the worm, rescues both its piRNA silencing and fertility phenotypes lends credence to the possible physiological importance of HDAC1 SUMOylation in this Alzheimer’s model. Taken together our worm studies and these studies in mammalian systems point to the likely importance of HDAC1 SUMOylation in the regulation of gene expression, and appear to validate the ‘SUMO-complementation by gene-fusion’ strategy employed here and previously by Tao et al., (2017).

### Parallels in the role of SUMOylation in Piwi silencing in flies and worms

Our proteomics studies reveal an intriguing stage-specific dynamic in the associations of HDA-1 and MEP-1, that is regulated in part by HDA-1 SUMOylation. In the adult germline, in the absence of HDA-1 SUMOylation, HDA-1 forms one or more complexes with a group of proteins including LIN-53/RbAP46/48, HPL-1, HPL-2, and MRG-1, but does not significantly engage a separate complex that includes MEP-1 and its co-factor LET-418/Mi2. Our findings suggest that HDA-1 SUMOylation promotes the association of these complexes in the adult germline. Moreover, although HDA-1 association with MEP-1 requires SUMOylation in the adult germline, in embryos their association is robust (Passannante et al., 2010; Unhavaithaya et al., 2002) and appears to be independent of HDA-1 SUMOylation (Kim et al., parallel).

Interestingly, in *Drosophila* Kc167 cells, a MEP-1-Mi-2 complex (called MEC) that lacks HDAC1 was reported (Kunert et al., 2009). Kc167 cells are derived from ovarian somatic cells and express components of the Piwi pathway (Vrettos et al., 2017) raising the question of whether HDAC1 SUMOylation might promote its association with the MEC complex in these cells. Interestingly, a recent paper showed that MEP-1 and the HDA-1 homolog RPD3 interact with PIWI and with Su(var)2-10 in fly ovarian somatic cells, where they function together to promote transposon silencing (Mugat et al., 2020). Thus it appears that in both worms and flies SUMO may promote assembly of a MEP-1/Mi-2/HDAC1 complex.

The worm ovary differs from these *Drosophila* systems in that the worm ortholog of Su(var)2-10 (GEI-17) is not required for piRNA-dependent silencing. Instead, the CCCH zinc-finger protein PIE-1 appears to bridge interactions between HDA-1 and the SUMO machinery (Kim et al., parallel). Appending SUMO as a fusion protein to HDA-1, HDA-1::SUMO, partially restored piRNA-reporter silencing when introduced into SUMO-pathway mutants (Figure 2C). These findings suggest that HDA-1 is a major target of SUMO in promoting Argonaute-dependent transcriptional silencing in the worm, and that E2 or E3 SUMOylation machinery is not required (or at least not continuously) once HDA-1 is modified. Another recent study found that the mouse Piwi homolog MIWI, engages NuRD complex components and DNA methyltransferases to establish de-novo silencing of transposons in the mouse testes, suggesting additional parallels to the findings from worms and flies (Zoch et al., 2020). However, studies in mice and Drosophila have not yet addressed the possibility that HDAC-1 is directly targeted by SUMOylation in promoting Piwi silencing.

The *mep-1* gene was previously implicated in regulating the transition from spermatogenesis to oogenesis in hermaphrodites (Belfiore et al., 2002), and our findings suggest that HDA-1 SUMOylation also promotes this function. We were surprised, however, to find that spermatogenesis targets are also regulated by the WAGO-pathway factor RDE-3 and by the Piwi Argonaute, PRG-1. Similar findings were also reported recently by Reed et al., (2020). These observations raise the interesting question of whether *C. elegans* germline Argonaute systems may function with SUMO and HDAC1 in regulating the switch from spermatogenesis to oogenesis in hermaphrodites.

## Materials and Methods

### C. elegans strains and genetics

All the strains in this study are derived from Bristol N2 and cultured on normal growth media (NGM) plates with OP50 and genetic analyses were performed essentially as described (Brenner et al., 1974). The strains used in this study are listed in Table S3.

### RNAi screen

RNAi screen was performed against all 337 genes in the chromatin subset of the C. elegans RNAi collection (Ahringer). RNAi of *smo-1* and *ubc-9* were added to the screen as chromatin regulators. Synchronous L1 worms of the reporter strain were plated on the IPTG plates with the corresponding RNAi food. Bacteria with Empty vector L4440 was used as negative control. The desilencing phenotype was scored when the worms on the control plates grew to young adult stage at 20 °C. In the first round of screen, 78 RNAi clones were scored positive. We performed second round of screen for these 78 positive ones and finally confirmed 29 of them including *smo-1* and *ubc-9* by requiring more than 5% of worms showing desilencing phenotype (Table S1).

### CRISPR/Cas9 genome editing

The Co-CRISPR strategy (Kim et al., 2014) and previously generated *mep-1* sgRNA, *smo-1* sgRNA (Kim et al., parallel) were used for *mep-1::gfp::degron/mep-1::mCherry::degron* and *3xflag::smo-1* strains, respectively. Other CRISPR lines were generated by Cas9 ribonucleoprotein (RNP) editing (Dokshin et al., 2018), or Cas12a (cpf1) RNP editing. Cas12a genome editing mixture containing Cas12a protein (0.5μl of 10 *μ*g/ *μ*l), two crRNAs (each 2.8μl of 0.2μg/μl), annealed PCR donor (4μg), *rol-6(su1006)* plasmid (2μl of 500ng/μl) was incubated at 37°C for 30min and 25°C for 1hr or O/N before injecting animals. For short insertions like FLAG, AID, HIS10 and point mutations, synthesized single strand DNAs were used as the donor; for long insertions like GFP, 2/3xFLAG-Degron, and SUMO, the annealed PCR products were used instead. The gRNA sequences were listed in Table S4.

### Generation of strains expressing SUMO conjugated hda-1

To prevent an additional conjugation of the fusion protein to other SUMO targets, we changed the tandem C-terminal glycines of SMO-1/SUMO to alanines (GG to AA) (Dorval and Fraser, 2006). The modified *smo-1* open reading frame was fused directly before the stop codon of the *hda-1, and hda-1(ne4747[KKRR])* gene at its endogenous locus by CRISPR genome editing as described above.

### Auxin treatment

For the AID system (Zhang et al., 2015), the *tir-1::mRubby* driven by a *sun-1* promoter and *eft-3* 3’ UTR mainly expressed in the germline were used. The degron tagged-L1 larval stage worms were plated on NGM plates with 100 μM auxin indole-3-acetic acid (IAA) (Alfa Aesar, A10556) unless otherwise stated, and kept in dark. Worms were collected at young adult stage for further analysis.

### Gonad fluorescent image

Gonads were dissected on glass slide (Thermo Fisher Scientific, 1256820) in M9 buffer, mounted in 2% paraformaldehyde (Electron Microscopy Science, Nm15710) in egg buffer (25mM HEPES pH 7.5, 118mM NaCl, 48mM KCl, 2 mM CaCl2, 2mM MgCl2), and directly imaged. Epi-fluorescence and differential interference contrast (DIC) microscopy were performed using an Axio Imager M2 Microscope (Zeiss). Images were captured with an ORCA-ER digital camera (Hamamatsu) and processed using Axiovision software (Zeiss).

### Immunofluorescence

Immunostaining of gonads was performed essentially as described (Kim et al., parallel). The primary antibodies (1:100) used were anti-acetyl-histone H3K9 antibody (Abcam, ab12179), anti-di-methyl-histone H3K9 antibody (Abcam, ab1220), and anti-tri-methyl histone H3K4 (Millipore, 07-473). The secondary antibodies (1:1000) used were goat anti-mouse IgG (H+L) Alexa Fluor 594 (Thermo Fisher Scientific, A11005), goat anti-mouse IgG (H+L) Alexa Fluor 488 (Thermo Fisher Scientific, A11001), and goat anti-rabbit IgG(H+L) Alexa Fluor 568 (Thermo Fisher Scientific, A11011), respectively. Epi-fluorescence and differential interference contrast (DIC) microscopy were performed using an Axio Imager M2 Microscope (Zeiss). Images were captured with an ORCA-ER digital camera (Hamamatsu) and processed using Axiovision software (Zeiss).

### Affinity chromatography of histidine-tagged SUMO

Synchronous adult worms (∼200,000) were used for Ni-NTA pull-down. HIS-tagged SUMO/SUMOylated proteins were purified as described in (Kim et al., parallel)

### Co-immunoprecipitation and western blotting

IP was performed as described (Kim et al., parallel). For the western blot analysis, protein samples to which NuPAGE LDS sample buffer (4x) (Thermo Fisher Scientific, NP0008) was added were loaded on precast NuPAGE Novex 4-12% Bis-Tris protein gel (Life Technologies, NP0321BOX) and transferred onto 0.2*μ*m nitrocellulose membrane (Bio-Rad, 1704158) using transblot turbo transfer system (Bio-Rad). The membrane was incubated with primary antibodies at 4 °C overnight and then with secondary antibodies for 1.5 hr at room temperature. Primary antibodies used were: anti-FLAG (1:1000) (Sigma-Aldrich, F1804), anti-GFP (1:1000) (GenScript, A01704) anti-MRG-1 (1:1000) (Novus Biologicals, 49130002), anti-HPL-1 (1:1000) (Novus Biologicals, 38620002), anti-HPL-2 (1:1000) (Novus Biologicals, 38630002), anti-LIN-53 (1:1000) (Novus Biologicals, 48710002), anti-HDA-1 (1:2500) (Novus Biologicals, 38660002), anti-LET-418 (1:1000) (Novus Biologicals, 48960002), anti-SMO-1 (Pelisch et al., 2014) (1:500) (purified from Hybridoma cell cultures, the Hybridoma cell line was gift from Ronald T. Hay, University of Dundee), anti-Ubiquitin (1:1000) (Abcam, ab7780), and anti-WAGO-9/HRDE-1(Ashe et al., 2012) (1:500) (gift from Eric A. Miska). Antibody binding was detected with secondary antibodies: goat anti-mouse (1:2500) (Thermo Fisher Scientific, 62-6520) and mouse anti-rabbit (1:3000) (Abcam, ab99697).

### Immunoprecipitation-Mass spectrometry (IP-MS)

Synchronous adult worms were used for IP experiments unless otherwise indicated. 1ml of frozen worm pellet was mixed with 1ml of IP lysis buffer (Kim et al., parallel) and 2ml of glass beads. Worm lysates were generated with FastPrep (MP Biomedicals) at a speed of 6 m/s for 20 s for four times. After clearing the lysates with centrifuge (12000 rpm, 30 mins at 4°C, twice), the GFP-tagged protein was pulled down with home-made anti-GFP/GBP beads (incubation for 1h at 4°C on a rotating shaker) and eluted with 1% SDS, 50 mM Tris, pH 8.0 at 95 °C for 10 minutes. Proteins were then precipitated with TCA and digested by Trypsin. LC-MS/MS analysis of the resulting peptides was conducted on a Q Exactive mass spectrometer (Thermo Fisher Scientific) coupled with an Easy-nLC1000 liquid chromatography system (Thermo Fisher Scientific). Peptides were loaded on a pre-column (75 μm ID, 6 cm long, packed with ODS-AQ 120 Å -10 μm beads from YMC Co., Ltd.) and separated on an analytical column (75 μm ID, 13 cm long, packed with Luna C18 1.8 μm 100 Å resin from Welch Materials) using an 78 min acetonitrile gradient from 0% to 30% (v/v) at a flow rate of 200 nl/min. Top 15 most intense precursor ions from each full scan (resolution 70,000) were isolated for HCD MS2 (resolution 17,500; NCE 27), with a dynamic exclusion time of 30 s. We excluded the precursors with unassigned charge states or charge states of 1+, 7+ or > 7+. Database searching was done by pFind 3.1 (http://pfind.ict.ac.cn/) against the *C. elegans* protein database (UniProt WS235). The filtering criteria were: 1% FDR at both the peptide level and the protein level; precursor mass tolerance, 20 ppm; fragment mass tolerance, 20 ppm; the peptide length, 6-100 amino acids. Spectra counts of HDA-1 in IP samples were used for normalization. Proteins either absent in N2 or with more than two fold of the spectra counts in the *hda-1::gfp* IP compared to those in N2, were shown in Table S2.

### Silver Staining

One third of samples from the IPMS experiment was loaded to a 4-12% SDS PAGE and the silver staining was carried out with ProteoSilver™ Plus Silver Stain Kit (Sigma). Each visible band was cut for trypsin digestion and MS identification, and the most abundant protein in each band was labeled in Figure 2A.

### RNAi inheritance

Worms at L1 stage were placed on IPTG plate (NGM plate with 1 mM IPTG and 100 μg/ml Ampicillin) seeded with *E. coli* strain HT115 transformed with either control vector L4440 or *gfp* RNAi plasmid. The worms were scored for OMA-1::GFP signal in the oocyte at gravid adult stage and transferred to regular NGM plates. The OMA-1::GFP was monitored at gravid adult stage for every generation till most worms recovered the expression of *oma-1::gfp*.

### Germline mortal assay

For each strain, 18 worms at L4 stage were singled and grown at 25°C. For each generation, the mothers were transferred to new plates every two days and their brood sizes were scored. One worm from each plate for every generation was randomly picked to continue score the brood size until the plate became totally sterile.

### Gonad mRNA-seq and analysis

For every mutant, about 100 gonads were dissected from the day-1 adult worms. We carefully cut at -1 position to make sure every gonad is similar. The RNA was extracted with Tri-reagent with a yield about 0.5 μg. Ribosomal RNA was depleted by RNase H digestion after being annealed with home-made anti-rRNA oligos for *C. elegans*. DNA was also removed by DNase treatment. RNA seq library was constructed with a KAPA RNA HyperPrep kit and sequenced at pair-end on a Nextseq 500 Sequencer with the illumina NextSeq 500/550 high output kit v2.5 (150 cycles).

For the quantification of the mRNA sequencing data, salmon was used to map the reads with the worm database WS268, and its output files were imported to DESeq2 in R. The differentially expressed genes were filtered by fold change more than 2 and adjusted p-value less than 0.05. The scatter plots were generated by the plot function in R.

### Small RNA cloning and data analysis

The small RNA cloning was conducted as the previous report (Shen et al., 2018). Worms were synchronized and collected at young adult stage. Small RNAs were enriched using a mirVana miRNA isolation kit (Invitrogen) from Trizol purified total RNA. Homemade PIR-1 was used to remove the d or triphosphate at the 5’ to generate 5’ monophosphorylated small RNA. Products were then ligated to 3’ adaptor (/5rApp/TGGAATTCTCGGGTGCCAAGG/3ddC/) by T4 RNA ligase 2(NEB) and 5’ adaptor (rGrUrUrCrArGrArGrUrUrCrUrArCrArGrUrCrCrGrArCrGrArUrCrNrNrNrCrGrArNrNrNrUrArCrNrNrN, N is for a random nucleotide) by T4 ligase 1(NEB) sequentially, followed by reverse transcription with RT primer (CCTTGGCACCCGAGAATTCCA) and SuperScript III (Invitrogen). PCR amplification was done with Q5 and primers with indexes (Forward: AATGATACGGCGACCACCGAGATCTACACGTTCAGAGTTCTACAGTCCGA,Reverse: CAAGCAGAAGACGGCATACGAGAT [6bases index] GTGACTGGAGTTCCTTGGCACCCGAGAATTCCA). PCR productions around 150 bp were separated by 12% SDS-PAGE, extracted with TE buffer (10 mM Tris-HCl, 0.1 mM EDTA, pH 8.0), and purified with isopropanol precipitation. Libraries were equally mixed and sequenced on a NextSeq 550 sequencer using the illumina NextSeq 500/550 high output kit v2.5 (75 cycles) with 75bp single-end sequencing. Adapters were trimmed by cutadapt and reads were mapped to a worm database (WS268) using Bowtie2. DESeq2 was used to normalized reads between samples.

### CHIP-seq

CHIP was carried out using the previous described protocol (Askjaer et al., 2014). Young adult worms were washed three times with M9 and once with PBS and then cross-linked with 1.1% formaldehyde in PBS with protease inhibitors for 10mins before being quenched with 125 mM glycine. Exchanging to CHIP Cell lysis buffer (20 mM Tris-Cl, 85 mM KCl, 0.5% NP40, pH8.0), DNA was fragmented with sonication (Bioruptor, High intensity, 30 sec on and 30 sec off for 45 cycles). Samples were first incubated with the corresponding antibodies overnight at 4 °C, and then with Magnetic beads, which were precleaned with 5% BSA in PBST (PBS+0.02% Tween-20), for 4 hours. After series of wash steps with buffer of different stringency, DNA was eluted with CHIP elution buffer (1% SDS, 250 mM NaCl, 10 mM Tris pH 8.0, 1 mM EDTA) at 65 °C for 15 mins twice. RNase A and Proteinase K treatments were carried out to remove the RNA and proteins. Reverse cross-linking was achieved by 65 °C overnight incubation and the DNA was purified with Zymo DNA clean kit (Cat # D5205). Libraries were prepared with NEBNext Ultra II DNA Library Prep Kit, equally mixed and sequenced on HiSeq X or NovaSeq 6000 with paired-end 150 bp sequencing. With a pair wised IP sample and its input, A CHIP-Seq Pipeline in DolphinNext platform built by the Bioinformatics core in UMass medical school was used to analyze the CHIP-seq data. Basically, it includes adapter removal (cutadapt), reads mapping (Bowtie2-align-s v2.2.3), duplicates removal (Picard-tools v1.131), peak calling (MACS2 v2.1.1.20160309), peak location and quantification (Bedtools v2.25.0, Quinlan and Hall 2010). The output bed files were used to generate Figures with IGV (v2.7.2) for Figure 4B. Background subtraction was applied with the intersect function of Bedtools for Figure S4.

## Acknowledgement

We thank members of Mello and Ambros labs for discussions. We thank M. Shirayama for sharing of strain and Miska and Hay labs for sharing antibodies. Onur Yukselen and Alper Kucukural from bioinformatics core of UMass Medical School provided support on CHIP-seq data analysis. C.C.M. is a Howard Hughes Medical Institute Investigator. This work was supported in part by NIH Grant GM 58800.

## Author Contributions

Conceptualization, H.K., Y.D., and C.C.M.; Investigation, H.K., Y.D., Y.Y., M.D., and C.C.M.; Writing-Original draft, H.K., Y.D., and C.C.M.; Writing-Review & Editing, H.K., Y.D., D.C., and C.C.M.; Supervision, C.C.M. and M.D.

