## Supplemental figures for "HDAC1 SUMOylation promotes Argonaute directed transcriptional silencing in *C. elegans*"

Figure S1 HDA-1(KKRR) caused increased levels of active chromatin H3K4me3

(A) Levels of H3K9ac levels was increased in Degron tagged *mep-1* strain without auxin exposure, compared to in WT (See figure 4A), which suggests leaky degradation of Degron fused MEP-1 protein (Yesbolatova et al., 2019).

(B and C) Active chromatin mark, H3K4me3 was increased in HAD-1(KKRR), compared to in WT. Immunofluorescence micrographs of anti-H3K4me3 and DAPI staining in adult gonad of WT in (B) and HDA-1(KKRR) in (C).

Figure S2 mRNA-seq changes and repeats.

(A) Number of up-(green) or down-(orange) regulated mRNAs (genes) with greater than 2-fold change and P value less than 0.05 from the indicated mutants' gonads. (B) Venn diagram showing the overlap of upregulated genes in *degron::hda-1*, *prg-1*, *rde-3*, *and wago-9*. The total number of the upregulated genes in each mutant is indicated in the parentheses.

(C to M) Scatter plot comparing of two independent repeats of the gonad mRNA sequencing results in the indicated strains. Two dashed lines outside indicate two-fold change.

Figure S3 mRNA-seq for transposons.

(A) Bar graphs showing the number of up-(green) or down-(orange) regulated transposons in the indicated mutants' gonads.

(B) Scatter plots of mRNA-seq data for the expression of transposons in the mutants. X axis is the log10 value of average reads counts in repeats for the control, Y axis is the log10 value of the reads count in mutants. Transposons with more than two-fold increase and p-value less than 0.05 are colored with red. The transposon's family names were labeled if available. WT was used as control for all the mutants except *degron::hda-1* and *ubc-9(ne4446[G56R]).* For *degron::hda-1*, the worms without auxin treatment were used as the control for 100 $\mu$M auxin treated, and *oma-1::gfp; gfp-csr-1*(AE) was used as the control for *ubc-9(ne4446[G56R]);oma-1::gfp; gfp-csr-1(AE)*. The dashed lines indicate two-fold change.

Figure S4 Anti-H3K9me2 CHIP-seq

Anti-H3K9me2 CHIP-seq in WT, *had-1(ne4747[KKRR])*, *degron::hda-1*, or *ubc-9(ne4446[G56R])* worms. Protein coding genes (A and B) and transposons (C and D) are shown. Reads counts (Y axis) are normalized by total reads based on those in WT.

Figure S5 mRNA-seq for protein coding genes.

Scatter plots of mRNA-seq data. X axis represents the log10 value of average reads in repeats for the control, Y axis for the mutants. WT was used as control for all the mutants except *degron::hda-1* and *ubc-9(ne4446[G56R]).* For *degron::hda-1*, the worms without auxin treatment were used as the control for worms with 100 $\mu$M auxin treatment, and *oma-1::gfp; gfp-csr-1*(AE) was used as the control for *ubc-9(ne4446[G56R]); oma-1::gfp; gfp-csr-1(AE)*. "Spermatogenic", "Oogenic" and "Neutral" genes were colored as indicated. A value of 0.1 was assigned to genes with missing reads. The dashed lines are for two-fold change

Figure S6 Small RNA-seq

(A)Scatter plot showing reads of anti-sense 22Gs targeting each gene in *hda-1(ne4747[KKRR])* versus those WT*.* The upregulated genes in *hda-1(ne4747[KKRR])* from mRNA-seq are indicated with red. Dashed lines represent two-fold change. (B) Ven diagram showing the overlap between upregulated genes in *hda-1(ne4747*[KKRR]) from mRNA-seq and genes with increased or decreased small RNAs in *hda-1(ne4747*[KKRR]). Total number of genes in each category is indicated in the parentheses.

Table S1 Summary of Chromatin RNAi screen

Table S2 HDA-1::GFP IP-MS

Table S3 List of *C. elegans* strains

Table S4 List of gRNA sequences

Table S5 RNA-seq data (deposited to Bioproject: PRJNA657279)

### Figure S1

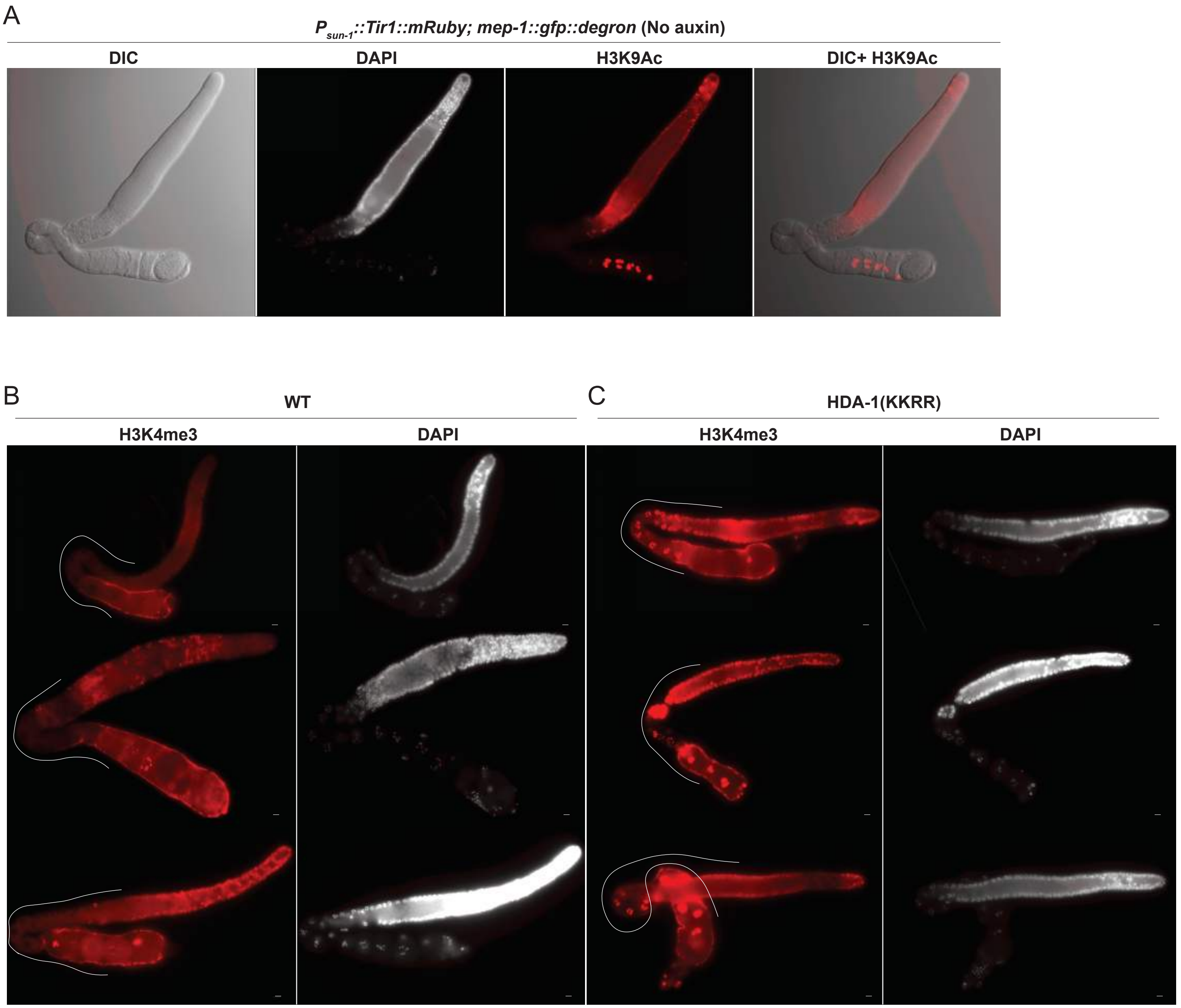
A
Psun-1::Tir1::mRuby; mep-1::gfp::degron (No auxin)
DIC
DAPI
H3K9Ac
DIC+ H3K9Ac
B
WT
H3K4me3
DAPI
C
HDA-1(KKRR)
H3K4me3
DAPI

### Figure S2

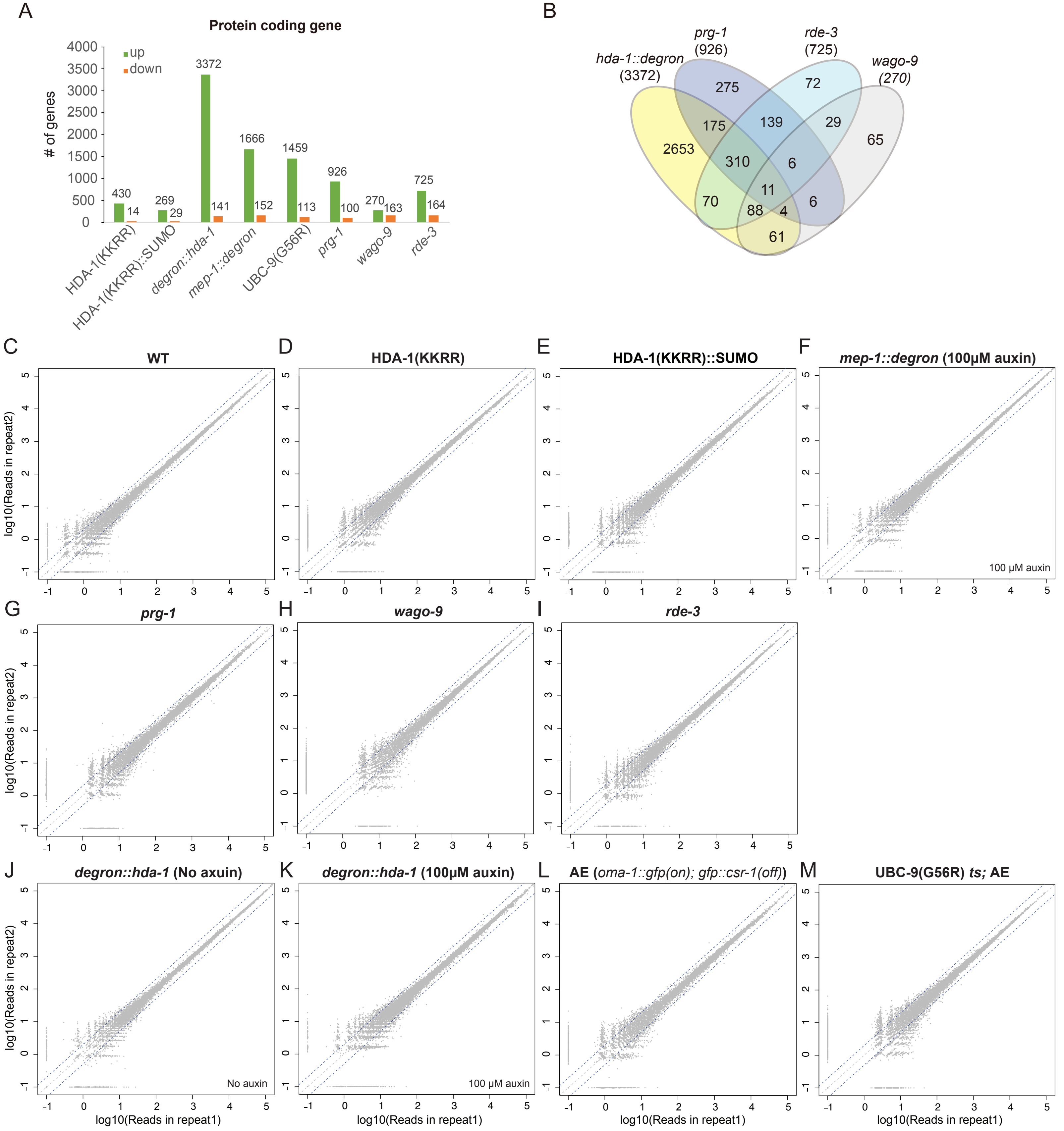
A
Protein coding gene
up
down
### of genes
HDA-1(KKRR)
HDA-1(KKRR)::SUMO
degron::hda-1
mep-1::degron
UBC-9(G56R)
prg-1
wago-9
rde-3
B
hda-1::degron (3372)
prg-1 (926)
rde-3 (725)
wago-9 (270)
C WT
D HDA-1(KKRR)
E HDA-1(KKRR)::SUMO
F mep-1::degron (100μM auxin)
100 μM auxin
G prg-1
H wago-9
I rde-3
J degron::hda-1 (No axuin)
No auxin
K degron::hda-1 (100μM auxin)
100 μM auxin
L AE (oma-1::gfp(on); gfp::csr-1(off))
M UBC-9(G56R) ts; AE
log10(Reads in repeat2)
log10(Reads in repeat1)

Figure S3

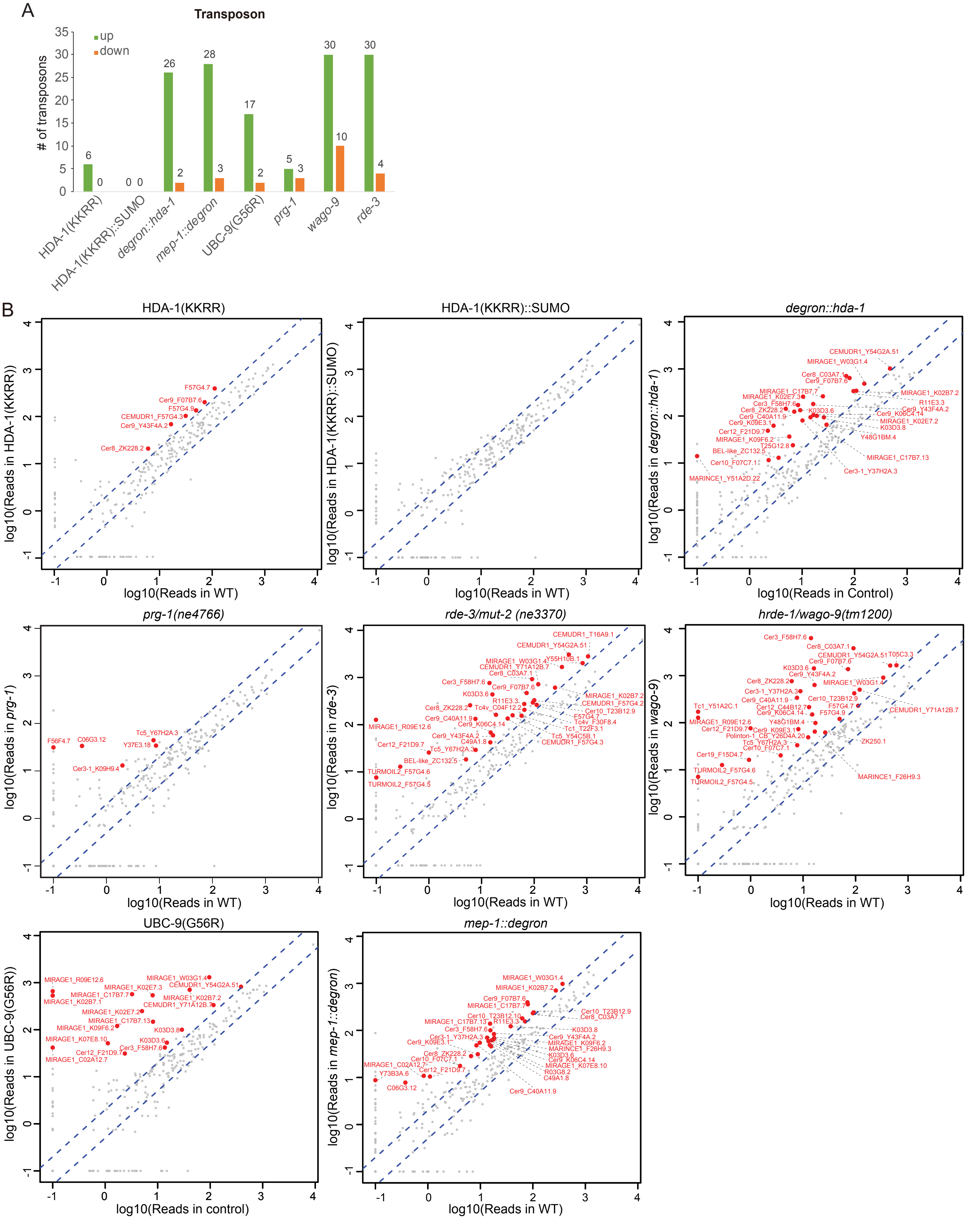
A
Transposon
up
down
### of transposons
HDA-1(KKRR)
HDA-1(KKRR)::SUMO
degron::hda-1
mep-1::degron
UBC-9(G56R)
prg-1
wago-9
rde-3
B
HDA-1(KKRR)
HDA-1(KKRR)::SUMO
degron::hda-1
prg-1(ne4766)
rde-3/mut-2 (ne3370)
hrde-1/wago-9(tm1200)
UBC-9(G56R)
mep-1::degron
log10(Reads in WT)
log10(Reads in Control)
log10(Reads in control)
log10(Reads in HDA-1(KKRR))
log10(Reads in HDA-1(KKRR)::SUMO)
log10(Reads in degron::hda-1)
log10(Reads in prg-1)
log10(Reads in rde-3)
log10(Reads in wago-9)
log10(Reads in UBC-9(G56R))
log10(Reads in mep-1::degron)

### Figure S4

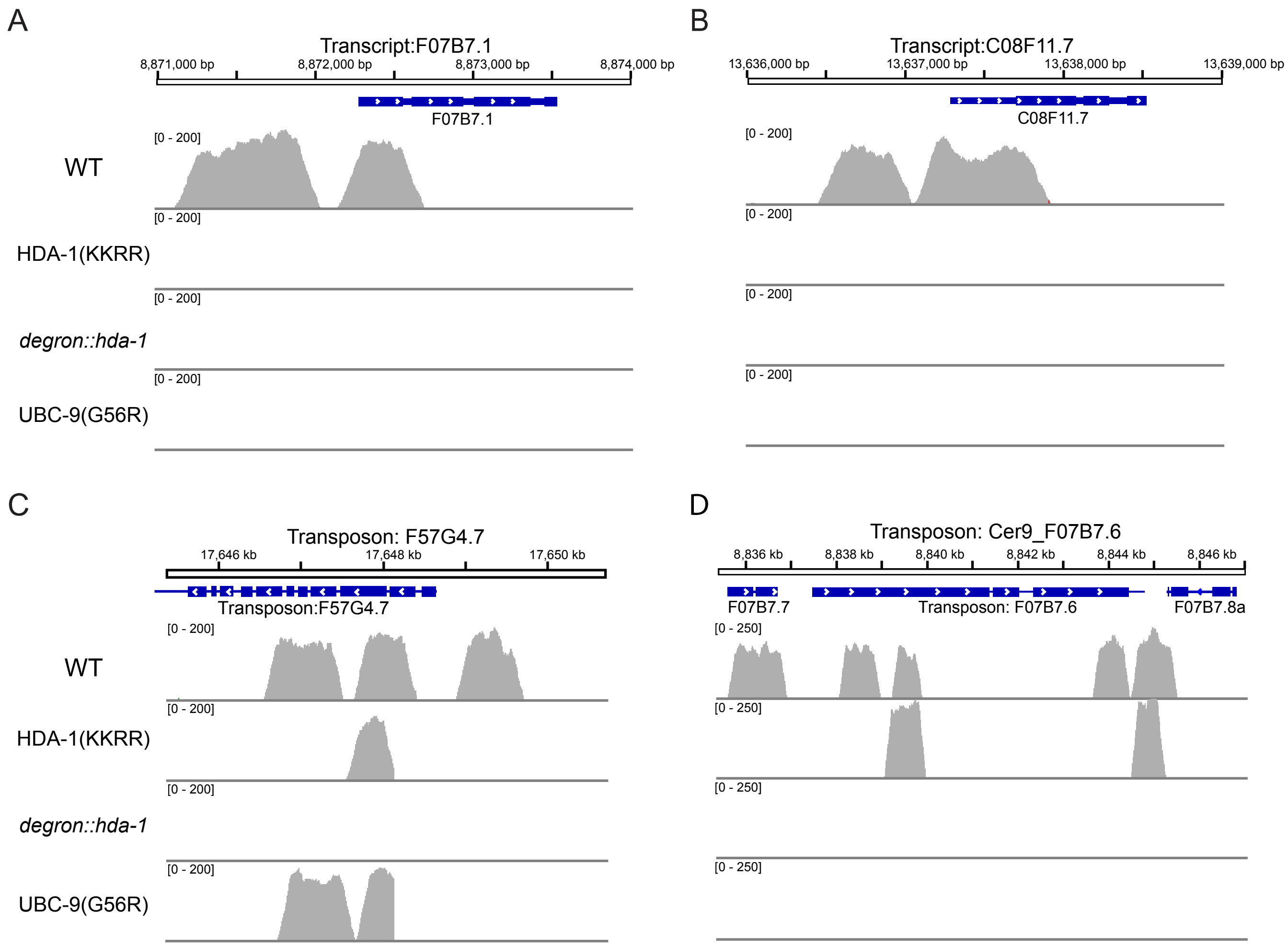

A
Transcript:F07B7.1
8,871,000 bp
8,872,000 bp
8,873,000 bp
8,874,000 bp
F07B7.1
[0 - 200]
WT
HDA-1(KKRR)
degron::hda-1
UBC-9(G56R)
B
Transcript:C08F11.7
13,636,000 bp
13,637,000 bp
13,638,000 bp
13,639,000 bp
C08F11.7
C
Transposon: F57G4.7
17,646 kb
17,648 kb
17,650 kb
Transposon:F57G4.7
D
Transposon: Cer9_F07B7.6
8,836 kb
8,838 kb
8,840 kb
8,842 kb
8,844 kb
8,846 kb
F07B7.7
Transposon: F07B7.6
F07B7.8a
[0 - 250]

Figure S5

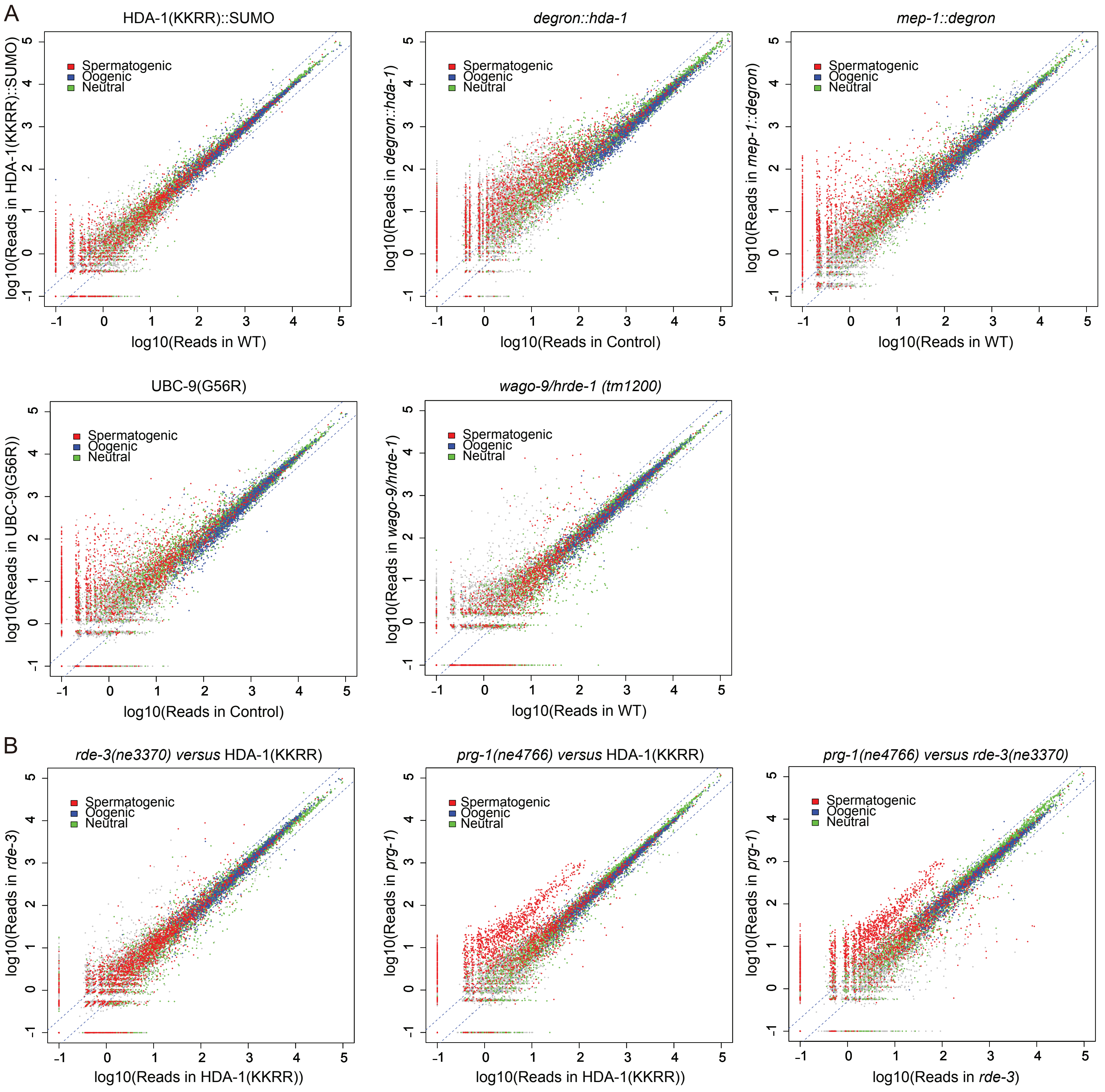
A
HDA-1(KKRR)::SUMO
log10(Reads in HDA-1(KKRR)::SUMO)
log10(Reads in WT)
Spermatogenic
Oogenic
Neutral
degron::hda-1
log10(Reads in degron::hda-1)
log10(Reads in Control)
mep-1::degron
log10(Reads in mep-1::degron)
log10(Reads in WT)
UBC-9(G56R)
log10(Reads in UBC-9(G56R))
log10(Reads in Control)
wago-9/hrde-1 (tm1200)
log10(Reads in wago-9/hrde-1)
log10(Reads in WT)
B
rde-3(ne3370) versus HDA-1(KKRR)
log10(Reads in rde-3)
log10(Reads in HDA-1(KKRR))
prg-1(ne4766) versus HDA-1(KKRR)
log10(Reads in prg-1)
log10(Reads in HDA-1(KKRR))
prg-1(ne4766) versus rde-3(ne3370)
log10(Reads in prg-1)
log10(Reads in rde-3)

Figure S6

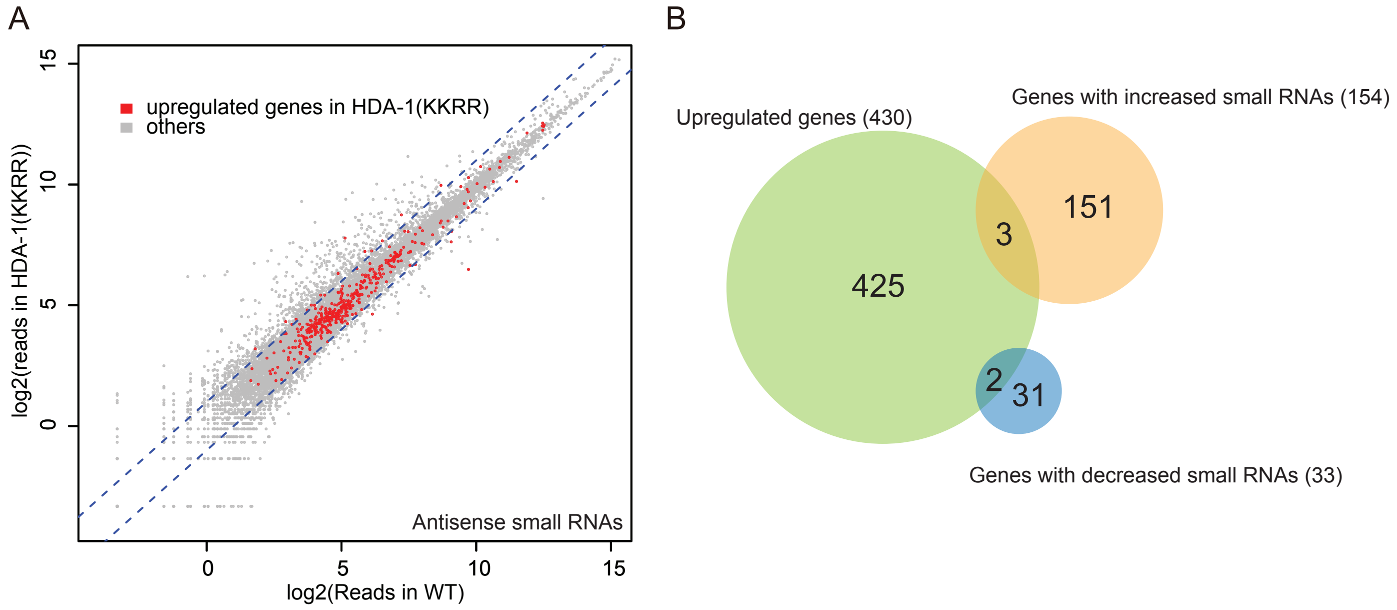
A
upregulated genes in HDA-1(KKRR)
others
log2(reads in HDA-1(KKRR))
Antisense small RNAs
log2(Reads in WT)
0
5
10
15
B
Upregulated genes (430)
Genes with increased small RNAs (154)
425
3
151
2
31
Genes with decreased small RNAs (33)
