## Supplemental Table 1 for "HDAC1 SUMOylation promotes Argonaute directed transcriptional silencing in *C. elegans*"

Table S1. Summary of Chromatin RNAi screen

| RNAi | Score | Number | Description |
| --- | --- | --- | --- |
| L4440 | 0% | N=20 | Empty vector |
| nrde-2 | 64% | N=28 | piRNA pathway |
| nrde-4 | 14% | N=28 | piRNA pathway |
| set-25 | 13% | N=23 | piRNA pathway |
| hda-1 | 14% | N=28 | NuRD complex |
| chd-3 | 6% | N=34 | NuRD complex |
| lin-53 | 12% | N=25 | NuRD complex |
| let-418 | 19% | N=26 | NuRD complex |
| sin-3 | 30% | N=27 | SIN-3 complex |
| dcp-66 | 8% | N=25 | HDA-1 interactor |
| lin-40 | 6% | N=34 | HDA-1 interactor |
| spr-5 | 8% | N=24 | CoREST |
| isw-1 | 11% | N=27 | SWI/SNIF complex |
| ssl-1 | 9% | N=23 | SWI/SNIF complex |
| set-33 | 17% | N=24 | histone methyltransferase |
| cbp-3 | 13% | N=24 | histone acetyltransferase |
| hda-2 | 9% | N=22 | histone deacetyltransferase |
| lin-61 | 5% | N=38 | Chromatin binding |
| mrg-1 | 12% | N=52 | Chromatin binding |
| lin-49 | 8% | N=24 | Bromodomain protein |
| trr-1 | 8% | N=25 | Transcription coregulator |
| din-1 | 10% | N=20 | transcriptional repressor |
| gmeb-3 | 11% | N=18 | transcription coactivator |
| taf-1 | 21% | N=19 | TATA-box binding |
| taf-5 | 10% | N=21 | TATA-box binding |
| air-2 | 56% | N=25 | Kinase |
| psr-1 | 7% | N=29 | arginine demethylase |
| acin-1 | 10% | N=30 | RNA binding |
| smo-1 | 28% | N=54 | SUMO |
| ubc-9 | 29% | N=58 | SUMO E2 |
