## Supplemental Table 3 for "HDAC1 SUMOylation promotes Argonaute directed transcriptional silencing in *C. elegans*"

Table S3. List of C. elegans strains

| Strain name | genotype | Method | Source |
| --- | --- | --- | --- |
| N2 | wild type |  | CGC |
| YY538 | *hrde-1(tm1200)* III |  | Buckley et al., 2012 |
| WM161 | *prg-1(tm872)* I |  | Yigit et al. 2006 |
| WM286 | *rde-3(ne3370)* I |  | Chen et al., 2005 |
| WM338 | *smo-1(ne4309*[3xFLAG::SMO-1]*)* I | CRISPR | This study |
| WM646 | *hda-1(ne4747*[HDA-1(K444R,K459R]*)* V | CRISPR | This study |
| WM647 | *hda-1(ne4748*[HDA-1(K444R,K459R)::HIS10::SUMO(GG to AA)]*)* V | CRISPR | This study |
| WM648 | *hda-1(ne4749*[HDA-1::HIS10::SUMO(GG to AA)]*)* V | CRISPR | This study |
| WM649 | *hda-1(ne4752*[3xFLAG::Degron::HDA-1]*)* V | CRISPR | This study |
| WM724 | neSi32[*Psun-1::Tir1::mRubby::eft-3 3'UTR*]; unc-119(ed9) III | MosSCI | This study |
| WM652 | neSi32 I; *mep-1(ne4629*[MEP-1::GFP::Degron]*)* IV | Cross | This study |
| WM653 | neSi22 [*oma-1::gfp (RNAa), cb-unc-119(+)*] II; neSi10 [*gfp::csr-1(RNAe), cb-unc-119(+)*] IV | Cross | Seth et al., 2018 |
| WM654 | neSi32 I; neSi22 II; neSi10 IV | Cross | This study |
| WM655 | *prg-1(tm872)* I;neSi22 II; neSi10 IV | Cross | This study |
| WM656 | neSi22 II; *hrde-1(ne4769[hrde-1 △]* III; neSi10 IV | Cross | This study |
| WM657 | neSi32 I;neSi22 II; neSi10 IV; *hda-1(ne4752*[3xFLAG::Degron::HDA-1]*)* V | CRISPR | This study |
| WM658 | neSi32 I;neSi22 II; neSi10 IV; *let-418(ne4757*[3xFLAG::Degron::LET-418]*)* V | CRISPR | This study |
| WM683 | neSi69[*Pwago-1::mep-1::flag::tev::gfp::wago-1UTR, cb-unc-119(+)*] II; unc-119(ed9) III | MosSCI | This study |
| WM684 | neSi22 II; neSi10, *mep-1(ne4755*[MEP-1::GFP::TEV::3xFLAG]*)* IV | Cross | This study |
| WM712 | *sin-3(tm1276)* I; neSi22 II; neSi10 IV | Cross | This study |
| WM713 | neSi22 II; *pie-1(ne4303*[PIE-1(K68R)]*)* III; neSi10 IV | CRISPR | This study |
| WM714 | neSi32 , *gei-17(ne4814*[2xFLAG::Degron::GEI-17]*)* I; neSi22 II; neSi10 IV | CRISPR | This study |
| WM660 | neSi22 II; neSi10 IV; *hda-1(ne4747*[HDA-1(K444R,K459R)]*)* V | Cross | This study |
| WM661 | neSi22 II; neSi10 IV; *hda-1(ne4748*[HDA-1(K444R,K459R)::HIS10::SUMO(GG to AA)]*)* V | Cross | This study |
| WM662 | *smo-1(ne4309*[3xFLAG::SMO-1]*)* I;neSi22 II; neSi10 IV | Cross | This study |
| WM663 | *smo-1(ne4309*[3xFLAG::SMO-1]*)* I;neSi22 II; neSi10 IV; *hda-1(ne4748*[HDA-1(K444R,K459R)::HIS10::SUMO(GG to AA)]*)* V | Cross | This study |
| WM664 | neSi22 II; neSi10, *ubc-9(ne4446*[ubc-9(G56R)]*)*IV | CRISPR | This study |
| WM666 | neSi22 II; neSi10, *ubc-9(ne4446*[ubc-9(G56R)]*)* IV; *hda-1(ne4748*[HDA-1(K444R,K459R)::HIS10::SUMO(GG to AA)]*)* V | Cross | This study |
| WM716 | neSi32 I, neSi22 II; neSi10, *mep-1(ne4629*[MEP-1::GFP::Degron]*)* IV | Cross | This study |
| WM667 | *smo-1(ne4346*[HIS10::SMO-1]*)* I | CRISPR | Kim et al, co-submitted |
| WM668 | *smo-1(ne4346*[HIS10::SMO-1]) I; *hda-1(ne4750*[HDA-1(K444R)]*)* V | CRISPR | This study |
| WM669 | *smo-1(ne4346*[HIS10::SMO-1]*)* I; *hda-1(ne4751*[HDA-1(K459R)]*)* V | CRISPR | This study |
| WM670 | smo-1(ne4346[HIS10::SMO-1] I; *hda-1(ne4747*[HDA-1(K444R,K459R)]*)* V | CRISPR | This study |
| WM671 | *mep-1(ne4380*[MEP-1::GFP::TEV::3xFLAG]*)* IV | CRISPR | Kim et al, co-submitted |
| WM672 | *mep-1(ne4380*[MEP-1::GFP::TEV::3xFLAG]*)* IV; *hda-1(ne4747*[HDA-1(K444R,K459R)]*)* V | Cross | This study |
| WM673 | *mep-1(ne4380*[MEP-1::GFP::TEV::3xFLAG]*)* IV; *hda-1(ne4748*[HDA-1(K444R,K459R)::HIS10::SUMO(GG to AA)]*)* V | Cross | This study |
| WM679 | *hda-1(ne4636*[HDA-1::GFP]*)* V | CRISPR | This study |
| WM680 | *hda-1(ne4754*[HDA-1(K444R, K459R)::GFP]*)* V | CRISPR | This study |
| WM682 | *prg-1(ne4766[prg-1△1181nt, null])* I | CRISPR | gift from Dr. Shirayama |
| WM685 | *hrde-1(ne4770*[GFP::TEV::2xFLAG::Degron::HRDE-1]*)* III | CRISPR | This study |
| WM710 | *gei-17(ne4822*[*gei-17 +3nt*, nonsense mutation]*)* I ; neSi22 II; neSi10 IV | CRISPR | This study |
| WM711 | *gei-17(ne4823*[*gei-17 △4nt*, nonsense mutation]*)* I ;neSi22 II; neSi10 IV | CRISPR | This study |
| WM715 | neSi32 I; *mrg-1(ne4824*[2xFLAG::Degron::MRG-1]*)* III; neSi22 II; neSi10 IV | CRISPR | This study |
