## Supplemental Table 4 for "HDAC1 SUMOylation promotes Argonaute directed transcriptional silencing in *C. elegans*"

Table S4. List of gRNA sequence

| guide | sequence | gene | usage |
| --- | --- | --- | --- |
| oYD274 | CATAATTTTGTCGAGCAAGT | hrde-1 | **3xflag**::hrde-1 |
| oYD218 | CATTCCGAAGCGAAACTTCT | hrde-1 |  |
| oYD426 | ccatcaaacATGAACTCAAA | hda-1 | **3xflag::degron**::hda-1 |
| oYD422 | TGAAGAAGATCCGTCTTTAG | let-418 | **3xflag::degron**::let-418 |
| HK_smo-1_sgRNA #5 | gagactcccgctataaacgA | smo-1 | **3xflag**::smo-1 |
| HK_pie-1_crRNA #2 | ATCTTGAGCGCTTCACGCTT | pie-1 | pie-1(**K68R**) |
| HK_ubc-9_crRNA #11 | AAGGATACGATTTGGGAAGG | ubc-9 | ubc-9(**G56R**) |
| HK_hda-1_crRNA1 | GCTCAGTTTGAGTCGGAAGG | hda-1 | hda-1(**K444R**) |
|  |  |  | hda-1(**K459R**) |
|  |  |  | hda-1(**K444R, K459R**) |
|  |  |  | hda-1::**gfp** |
| HK_hda-1_cpf1_crRNA1 | CTCTGTCTTCTGACGCTTTTC | hda-1 | hda-1(WT)::**his10::SUMO(GG to AA)** |
| HK_hda-1_cpf1_crRNA2 | GTGTTTTACTCTGTCTTCTGA | hda-1 |  |
| HK_hda-1_cpf1_crRNA3 | CTCCGTACGCTGACGCTTTT | hda-1(KKRR) | hda-1(K444R,K459R):**his10:::SUMO(GG to AA)** |
| HK_hda-1_cpf1_crRNA4 | GTGTTTTTACTCCGTACGCT | hda-1(KKRR) | hda-1**(**K444R,K459R**)**::**gfp** |
| HK_mrg-1_crRNA 1 | TTCCTTTGAAGACATctga | mrg-1 | **2xflag::degron**::mrg-1 |
| HK_gei-17_crRNA1 | GGTATTTGCCATTGATTATT | gei-17 | **2xflag::degron**::gei-17 |
| HK_gei-17_crRNA2 | atATGTTACCGAATAATCAA |  | gei-17 nonsense mutaion |
| HK_mep-1_sgRNA #4 | GCGCAAAAGAAGGAAGACGG | mep-1 | mep-1::**mCherry::degron** |
|  |  |  | mep-1::**gfp::degron** |
